# Histo-LCM-Hi-C reveals the 3D chromatin conformation of spatially localized rare cells in tissues at high resolution

**DOI:** 10.1101/2025.06.10.658807

**Authors:** Yixin Liu, Min Chen, Xin Liu, Zeqian Xu, Xinhui Li, Yan Guo, Daniel M. Czajkowsky, Zhifeng Shao

## Abstract

It is now well established that an understanding of the chromatin structure is essential to delineate the mechanisms underlying genomic processes. However, while methods to obtain this information from cells *in vitro* are widely available, there is presently a significant lack of techniques that can acquire this data from cells in the tissue. Such a capability is critical to determine the dependence of the local tissue environment on cell functioning. Further, this ability is particularly necessary for cells that are a significant minority of the total tissue population, which are often obscured in data dominated by more abundant tissue cells. Here we have developed Histological Laser Capture Microdissection Hi-C (Histo-LCM-Hi-C) to enable the characterization of chromatin architecture of phenotype-defined, spatially localized cells within intact tissue sections from as few as about 300 cells. We demonstrate the effectiveness of this approach with the generation of the first 3D Hi-C map of the tissue-resident macrophages of the liver, the Kupffer cells (KC), which are a minor cell population in the normal liver. As expected, owing to their relative rarity, these KC maps are significantly different from those obtained from whole liver, revealing distant contacts between putative enhancers and genes involved in key KC functions as well as significant differences with that of *in vitro* induced bone-marrow derived macrophages. We anticipate that this method will prove to be an indispensable technique in the growing repertoire of methodologies used for the characterization of the genomic properties of cells within their native environment.

## Introduction

It has long been recognized that organs are composed of cells with a wide range of phenotypes whose functioning is tightly coordinated to ensure the proper functioning of the whole organ. A critical aspect of this coordination is the precise spatial organization of the cells within the tissue: the interactions of the cells with their local milieu profoundly influence gene expression and cellular function [1–3]. Indeed, the recent proliferation of spatial -omics technologies has revealed a tremendous amount of information that has significantly advanced our understanding of cellular functioning within native tissues [4–6]. However, among these spatial -omics methods, there is a notable lack of techniques that are capable of providing genome-wide views of chromatin architecture. This is in spite of the fact that it is now clear that the 3D confirmation of the genome plays a fundamental role in many genomic processes, most notably transcription [7, 8]. Indeed, an understanding of chromosome structure is now recognized as ultimately essential to identify contacts between enhancers and promoters that play a pivotal role in the regulation of gene expression [9].

To date, one of the most common methods to study chromatin architecture *in vitro* is high-throughput chromosome conformation capture (Hi-C) [10]. With this method, structures over a range of genomic scales have been identified, from point-to-point contacts that define loops [11], to more extended regions of enhanced contacts that span hundreds to thousands of base pairs called topologically-associated domains (TADs) [12], to much longer-range contacts of broad regions that define compartments [13]. The Hi-C method employs a series of enzymatic steps that are highly effective for cultured cells but are far less effective when applied to intact blocks of tissue, owing to the reduced diffusion of enzymes within the dense matrix of the tissue. Further, the Hi-C maps that could be obtained from such samples would be averages over the entire population of the cells in the tissue, regardless of their positioning or phenotype. Indeed, to our knowledge, there is presently no established method to obtain Hi-C maps from spatially defined cells within intact native tissue.

With this in mind, we recognized that Laser Capture Microdissection (LCM) is capable of isolating specific cells directly from tissue sections [14, 15]. With LCM and an appropriate marker, even rare cells can be easily identified and selected for analysis, ensuring a direct and reliable characterization of these cells. We note that characterizing rare cells in tissues is presently a well-recognized challenge for many of the spatial -omics methods developed to date [4]. Here, we present a novel protocol to acquire high quality Hi-C data from as few as 300 cells using LCM, with the cells specifically identified using standard histological-based staining techniques (Histo-LCM-Hi-C). We demonstrate the power of this approach by generating the first Hi-C maps of Kupffer cells directly isolated from mouse liver sections. These maps reveal previously unknown details of enhancer-promoter contacts that are specific for KC cells and are associated with genes that have pivotal roles for KC functioning. We anticipate that this methodology will find broad application among the growing arsenal of spatial -omics methods used to study the functioning of native tissues.

## Materials and methods

### Preparation of the mouse tissue sample

The samples were obtained from age- and gender-matched mice, namely four and two 12- to 13-week-old male C57BL/6 mice (Jie Si Jie Laboratory Animals, Shanghai, China) for the liver and brain samples, respectively. Mice were housed under ad libitum feeding conditions and maintained on a reversed light–dark cycle. Mice were euthanized by cervical dislocation 6 hours after turning off the lights. The liver was then freshly dissected, washed with ice-cold phosphate-buffered saline (PBS), and cut into small blocks. For the optimized method, the tissue blocks were fixed in 1% paraformaldehyde at 17 °C for 4 hours, followed by quenching with 0.125 M glycine for 15 minutes at room temperature. The blocks were stored at 4 °C.

### *In situ* Hi-C in tissue sections

The tissue blocks were placed in ice-cold PBS and cut into 8 to 10 µm thick sections using a vibratome (speed: 0.50 mm/s, amplitude: 0.75). Hi-C was then performed following a published method [16] with modifications. For the optimized method, the sections were incubated in lysis buffer [10 mM Tris-HCl, pH 8.0; 10 mM NaCl; 0.2% Tween-20; 1% Protease Inhibitor Cocktail (Catalog No. P8340, Sigma-Aldrich, St. Louis, MO)] on ice for 20 minutes, followed by washing with NEBuffer 2 (Catalog No. B7002S, NEB, Ipswich, MA). The sections were then treated with SDS (final concentration: 1.2%) at 65 °C for 1 hour, followed by incubation with Triton X-100 (final concentration: 1%) at 37 °C for 15 minutes. The samples were digested overnight at 37 °C with MboI restriction enzyme (Catalog No. R0147, 5 U/mL, NEB, Ipswich, MA), end-repaired with Klenow enzyme (Catalog No. M0210, NEB, Ipswich, MA) at 37 °C for 2 hours, and ligated using T4 ligase (Catalog No. M0202, NEB, Ipswich, MA) at 25 °C for 8 hours. Finally, the samples were treated with Exonuclease III (Catalog No. M0206, 100 U/mL, NEB, Ipswich, MA) for 5 minutes at 37 °C and stored at -80 °C until further processing. Analysis of the gel data was performed using ImageJ (version 1.54p; https://imagej.net/ij/).

### Labeling of KCs

The sections were washed three times with ddH2O for 5 minutes and then incubated with 3% hydrogen peroxide at room temperature for 10 minutes. The sections were blocked with 5% BSA, followed by washing with PBS. Next, the sections were incubated with the primary antibody, anti-rabbit F4/80 antibody (Catalog No. 70076s, Cell Signaling Technology, Danvers, MA), at 4 °C overnight. Afterward, the sections were washed with PBST (1 × PBS with 0.1% Tween). The sections were then incubated with goat anti-rabbit IgG H&L (HRP) [Immunohistochemistry Kit (Catalog No. G1215-200T, Servicebio, China) for 2 hours at room temperature. The sections were washed three times with PBS, 5 minutes each time. Then, 50 to 100 µL of DAB working solution was added and incubated at room temperature for several minutes, until the desired color developed under the microscope. Following this, the sections were washed with ddH2O to terminate the color development. Finally, the sections were counterstained with hematoxylin for about 1 to 2 minutes and washed with ddH2O for 2 minutes. If the tissue sections were used for LCM selection, the counterstaining step was omitted.

### Laser Capture Microdissection

Following histo-labeling, the sections were placed on a polyethylene terephthalate (PET)-membrane-covered slide (Catalog No. 415190-9051-000, Carl Zeiss, Jena, Germany), and the KCs were micro-dissected at the single nuclear region level using the Zeiss PALM Micro Beam LCM system (Zeiss Microimaging, Munich, Germany) housed in a Plexiglas housing. In short, the perimeter of the chosen cell was first finely cut with the focused laser beam whose resolution is less than 1 µm. This resolution ensures that only the chosen cell and, in particular, its nucleus is collected, and not that of the adjacent, neighboring cells. After this cutting, the isolated region is then ejected into the adhesive cap of the collection tubes (Catalog No.415190-9191-000, Carl Zeiss, Jena, Germany) using a pulse from a UV laser. We continued this process for the KCs, collecting material one cell at a time until sufficient materials were accumulated. To avoid external contamination, the LCM system was housed in a dedicated plexiglass chamber, which isolates all sorting, dissection, and ejection steps from the external environment. We also inspected the adhesive cap of the collection tubes under phase-contrast microscopy to confirm the absence of debris or foreign particulates, and subsequently placed the tube in a Biological Safety Cabinet (1300 Series A2, Thermo Fisher Scientific) to perform the subsequent experimental steps.

### DNA extraction and Hi-C library preparation

The collected sample was incubated with proteinase K (1 mg/mL) and SDS (1%) at 65 °C for 3 hours, followed by the addition of sodium chloride to a final concentration of 500 μM, and incubated at 65 °C overnight, with all steps from the opening of the adhesive cap to buffer addition performed on a dedicated workbench to prevent environmental contamination. The DNA was captured using 1.8 × Agencourt AMPure XP beads (Catalog No. A63881, Beckman, Brea, CA). The DNA was then resuspended in 10 mM Tris-HCl, and Tn5 enzyme [TruePrep DNA Library Prep Kit V2 for Illumina, (Catalog No. TD502/TD503, Vazyme, China)] was added to fragment the DNA and attach adaptors. Biotin-labeled chimeric molecules were captured by incubating with Dynabeads® MyOne™ Streptavidin C1 beads (Catalog No.65001, Thermo Fisher Scientific, Baltics UAB) at room temperature for 2 hours. After incubation, the beads were washed with 1 × Tween wash buffer (0.05% Tween 20, 1 M NaCl, 0.5 mM EDTA-NaOH, 5 mM Tris-HCl, pH 8) on a Thermomixer at 55 °C for 5 minutes, repeated four times. After washing, the DNA on the beads was amplified using the TruePrep™ DNA Library Prep Kit V2 (Catalog No. TD503, Vazyme, China). Finally, 0.55 × and 0.75 × Agencourt AMPure XP beads were used to select DNA fragments of the appropriate size for sequencing. All libraries in this study were sequenced on the NovaSeq 6000 platform. We note that standardized protocols were implemented for tissue fixation, LCM, Hi-C library preparation, and downstream data analysis for all samples.

### Hi-C data analysis

Hi-C reads were iteratively mapped to the mm10 genome using Bowtie2 (v2.1.0) with a step size of 25 bp. Reads that mapped to multiple genomic locations or had low mapping quality (MAPQ < 30) were removed. PCR duplicates were further removed. Read pairs identified as originating from the same fragment (i.e., pairs mapped to the same restriction fragment), nonspecific ligations (fragments with sizes above 600 bp), dangling ends (inward-facing read pairs less than 1 kbp apart), and self-ligations (outward-facing read pairs less than 25 kbp apart) were excluded. The valid pairwise reads were retained for each cell. Juicer tools (v1.19.02) were used to create the .hic file and perform KR normalization. The similarity between Hi-C data sets was calculated using HiCRep [17]. Compartment analysis was performed using Juicer tools at 100 kb bin size (except where noted), with the eigenvector corrected based on gene density. Regions with values greater than 0 were considered A compartment, and those with values less than 0 were considered B compartment. TADs were identified using TopDom [18] at 40 kb bin size, with a window size of 5; only regions with p-values < 0.05 were retained. For comparisons between samples, TAD borders within one 40 kb bin of each other were considered as co-localized. The valid pairwise reads per cell were calculated by dividing the total valid pairwise reads of one library by the number of cells in the library, the latter of which was determined based on the quantity of DNA. FitHiC2 [19] was used to identify significant Hi-C contacts. All bins with very low contact frequencies (namely, bins that contact fewer than 3% of the other bins) were excluded from this analysis. This software builds its empirical null model based on the input data and so does not require a strict sequencing depth or resolution (ref 19). For all analysis, based on our sequencing depth, we used a bin size of 20 kb to ensure greatest confidence in obtained results. For calculations of contacts with super-enhancers, we used a threshold q value of 0.005 to select for the most significant contacts with these functional elements. We used Mustache [20] to identify the loops in the Hi-C data with the sparsityThreshold value set at 0.88 and a p-value threshold of 0.05. We also used diffHic [21] to identify local differences in the Hi-C contact maps with a 40 kb bin size, with |logFC| greater than 0.4 and p-value of 0.05 as thresholds. The published AML12 cell Hi-C data, brain Hi-C data, and BMDM Hi-C data can be found at GEO with the accession number GSE136306, GSE34587, and GSE115524, respectively. The published liver Hi-C data are available at GEO with accession numbers GSE65126, GSE104129, and GSE58752.

### Identification of super enhancers

Super-enhancers (SEs) were identified using the Rank Ordering of Super-Enhancers (ROSE) algorithm [22, 23]. This approach was employed to delineate transcriptional regulatory elements throughout the genome, specifically focusing on regions located at least 3 kilobases away from transcription start sites (TSSs). To achieve this, adjacent H3K27ac ChIP-seq signals were integrated using the parameter “-t 3000.” Subsequently, the highest-ranked regions were classified as super-enhancers in the KC cells. The published H3K27ac ChIP-seq data for KCs is available in GSE216164.

### RNA-seq data analysis

Raw reads were processed with Trimmomatic (v0.35) to remove low-quality reads using the parameters “SLIDINGWINDOW:3:20 LEADING:20 TRAILING:20 MINLEN:30”. rRNA reads were removed using SortMeRNA. FastQC (v0.11.5) was used to check the quality of the clean reads. The clean reads were then mapped to the mm10 genome using HISAT2 (v2.0.5). Gene expression levels were calculated as transcripts per million (TPM) using StringTie (v1.3.3), with only mRNAs included. The Mus_musculus.GRCm38.96.chr_mRNA.gtf file was used and downloaded from http://ensemblgenomes.org/. TSS information was obtained from the same .gtf file. GO analysis was performed using ShinyGo [24] and Metascape [25]. The published KCs RNA-seq data and liver (hepatocyte) RNA-seq data are available at GEO with the accession numbers GSE186327 and GSE150835, respectively.

## Results

### Overview of Histo-LCM Hi-C

The workflow of this approach is outlined schematically in **Figure 1**. Briefly, dissected tissues are first crosslinked with paraformaldehyde and then sliced into tissue sections with a vibratome. The sections are then suspended in solution and the digestion and ligation steps of *in situ* Hi-C are performed. After the completion of these steps, the treated sections are deposited onto glass slides, histo-chemically stained, and then transferred to the specific slides used for LCM, followed by LCM to isolate specific cells from the stained sections. Finally, the Hi-C library is prepared from the isolated materials using slightly altered methods from published protocols [26] and the DNA is sequenced.

**Figure 1.**
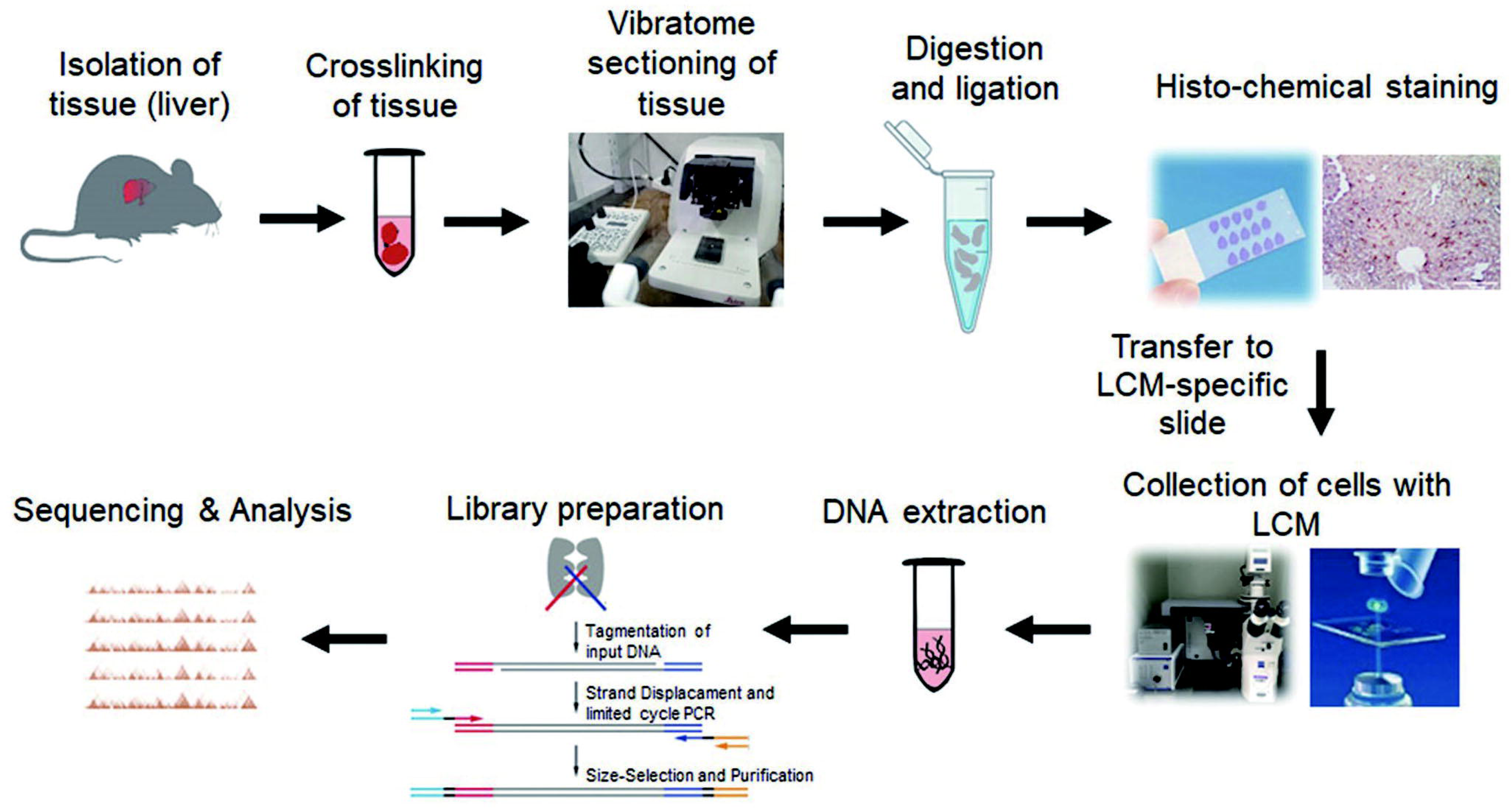
Overview of the Histo-LCM-Hi-C method.

### Optimization of the Hi-C protocol for tissue sections

As described in the Introduction, performing the enzymatic reactions of Hi-C with an intact whole tissue is generally challenging owing to the reduced diffusion of enzymes through the densely crosslinked tissue. Consequently, most prior Hi-C experiments involving tissues have dissociated the constituent cells into single cells or nuclei suspensions and performed the Hi-C procedure with these suspensions. However, such approaches prohibit any opportunity to understand the spatial dependence of the properties of the cells within the tissue. Moreover, simply isolating the cells from the tissue can also lead to alterations in chromatin structure [27]. Therefore, we sought to develop a Hi-C based method to directly characterize defined cell types within the tissue.

To this end, we first focused on identifying effective conditions for crosslinking the sample, since this step is critical to not only ensure a native chromatin structure but also allow efficient restriction/ligation during the Hi-C protocol in the tissue section. To identify these conditions, we used micrococcal nuclease (MNase) as a general means of characterizing chromatin accessibility to enzymatic digestion [28]. In particular, we used the extent of digestion to mono-nucleosome-spaced bands as an indicator of a practically useful level of crosslinking. We note that it is essential to achieve a uniform and sufficient crosslinking across the sample, which is generally much more challenging for a tissue/organ than for cultured or isolated cells. With blocks of mouse liver tissue, we found that typical tissue crosslinking conditions (4% paraformaldehyde for several days at 4 °C [29]) resulted in samples that were too heavily crosslinked for MNase digestion. In contrast, milder crosslinking conditions (1% paraformaldehyde for 10 min at room temperature, as used in standard Hi-C) produced smeared bands after MNase digestion (Figure S1A), indicating insufficient crosslinking. Thus, we examined the effectiveness of fixation for different times (2 hr, 3 hr or 4 hr) and also at different temperatures (4 °C, 17 °C and room temperature), monitoring the level of MNase digestion and tissue mechanical strength. In the end, we found that short fixation times (< 3 hr) were not effective, as this resulted in overly soft tissues that could not be efficiently sectioned. Moreover, low temperatures failed to preserve chromatin integrity. Hence, we settled on 1% paraformaldehyde for 4 hours at 17 °C, which yielded clear nucleosome-sized MNase digestion bands, thus indicating well-crosslinked samples (Figure S1B; Methods). We have also compared the results obtained using this fixation protocol with cultured AML-12 cells in relation to those obtained with the conventional fixation protocol, and found excellent agreement (Figure S1C).

With these conditions, we next focused on identifying the optimal section thickness for the Hi-C reactions. Since the size of a typical nucleus is in the range of 8 to 15 μm, we first examined sections of thickness between 10 μm to 50 μm for the extent to which the restriction enzyme used in Hi-C protocol (MboI) can digest the genomic chromatin. We performed this digestion, and all Hi-C steps, with freely floating sections so the enzymes could access the sections equally from all sides. However, we were surprised to find that the restriction digestion reaction was poor for thicknesses greater than 10 µm, even though the aforementioned MNase digestion assay was effective at these thicknesses (Figure S2A, S2B and S2C). We speculate that this reduced effectiveness was owing to a reduced diffusion of the MboI enzyme over MNase as a result of its larger size [30, 31]. Nonetheless, for this method, we used section thicknesses of 10 µm.

We next optimized the key parameters of the Hi-C procedure. We found that increasing the SDS treatment time improved digestion efficiency (Figure S2D; Methods). In addition, extending the times for end-repair and T4 ligase treatment significantly increased the ligation efficiency, yielding a sharp high-molecular weight band in the agarose gel as expected (compare Figure S3A and S3B). The library produced with these conditions was also of high quality (Figure S4A). Finally, we confirmed that these experimental conditions were also effective with a different tissue, namely mouse brain cortex (Figure S3C).

With these technical issues optimized, we examined the overall effectiveness of our Hi-C protocol with mouse liver sections and brain sections. As shown in Figure S4A& Figure S4B and **Table 1**, the sequenced libraries prepared with our protocol were of high quality and comparable to that of cultured AML 12 mouse hepatocyte cells obtained with the standard Hi-C protocol [16]. In addition, the maps obtained with our method were highly reproducible, exhibiting a stratum-adjusted correlation coefficient [17] (SCC, bin size 500 kb) of 0.98 between biological replicates **(Figure 2A**). By contrast, the SCC between the liver and brain sections was notably lower, at 0.64 (Figure 2A). We note that the SCC between the liver and AML 12 cells is also low (SCC of 0.55), suggesting that this cell line may not, in fact, accurately reflect the native hepatocytes in the tissue (which comprise ∼65 to 70% of the cells in the liver [32]).

**Figure 2.**
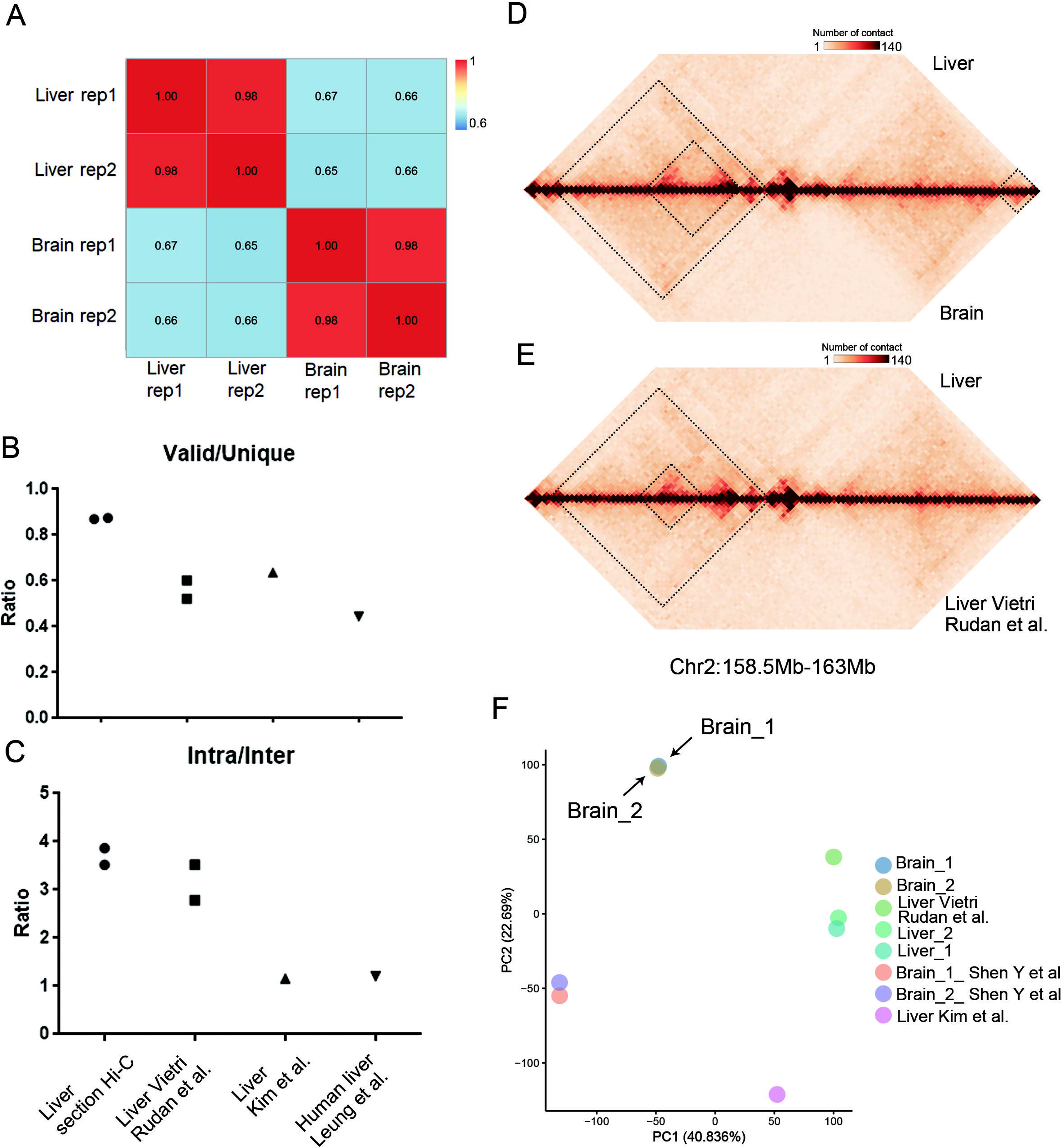
Comparison of the Hi-C data from brain and liver tissue sections with each other and existing datasets. **A.** Correlation results between the tissue section Hi-C data of liver and brain replicates. **B.** The proportion of valid pairwise reads in the liver tissue section Hi-C compared with the published mouse liver cell standard Hi-C data. **C.** The ratio of intra-chromosomal pairwise reads to inter-chromosomal pairwise reads in liver tissue section Hi-C as well as that for the published mouse liver cell standard Hi-C data. **D.** and **E.** Visualization of contact maps generated from tissue section Hi-C of liver and published mouse liver cell standard Hi-C data (**D**) as well as tissue section Hi-C of brain (**E**). Resolution: 40 kb. **F.** Comparison of the similarity of the data obtained from the indicated samples based on their corrected PC1 values (where PC1 values are typically used in compartment analysis) [33, 34, 36].

**Table 1.** Hi-C data from tissue sections and the cell line, AML 12.

|  | Brain-1 | Brain-2 | Liver-1 | Liver-2 | AML12-1 | AML12-2 |
| --- | --- | --- | --- | --- | --- | --- |
| Raw Reads Count | 160,848,154 | 175,003,865 | 53,552,169 | 194,377,862 | 213,721,058 | 195,013,809 |
| MAPQ≥30 Count | 99,040,570 | 107,767,081 | 38,695,055 | 140,320,344 | 141,943,465 | 126,168,943 |
| Duplicate Removed | 76,168,576 | 92,854,661 | 27,893,906 | 100,608,487 | 101,532,358 | 108,059,298 |
| Same Fragment | 10,625,104 | 8,862,075 | 1,066,341 | 3,463,072 | 19,687,823 | 12,407,978 |
| Nonspecific | 12,644,995 | 12,021,881 | 2,235,063 | 7,361,541 | 21,472,368 | 14,688,103 |
| Dangling | 11,394,617 | 16,315,089 | 1,144,714 | 4,271,417 | 14,713,256 | 14,998,043 |
| Self- Ligations | 628,856 | 754,687 | 117,354 | 451,430 | 836,685 | 1,026,657 |
| Valid Pairs | 50,631,175 | 63,200,700 | 24,253,195 | 88,019,801 | 63,402,300 | 76,437,463 |
| Cis-interaction | 40,822,535 | 52,653,103 | 18,883,608 | 69,929,164 | 42,289,461 | 52,565,598 |
| Trans-interaction | 9,808,640 | 10,547,597 | 5,369,587 | 18,090,637 | 21,112,839 | 23,871,865 |

We also found that the quality of our liver Hi-C data was comparable to or better than that of previously published data from mouse liver cells [33–35] (**Figure 2B** and **2C**) and AML 12 cells, the latter two of which were obtained using standard Hi-C (Figure S5). In addition, over large genomic distances (namely, at a bin size of 500 kb), our liver section data were highly correlated with this previously published data (SCC of 0.95 and 0.92). However, we found that, over small distances (at a bin size of 40 kb), this correlation was lower (SCC of 0.90 and 0.77), suggesting differences in structure at the TAD/sub-TAD scale (Figure S6A and S6B). Such differences between our liver section data and previously published work were also apparent by direct inspection of the Hi-C contact maps (**Figure 2D)**. We note though that such differences were less frequent than those between the liver and brain samples (**Figure 2E**). Examination of these maps at the level of compartments (obtained at a bin size of 100 kb) also showed a similar trend, with clear differences between the liver section and the published liver work (**Figure 2F**). A similar finding was also observed for the brain tissue section Hi-C data and published brain Hi-C data [36] (SCC of 0.89 and 0.81 for bin sizes of 500 kb and 40 kb, respectively) (Figure S6A and S6B; Figure 2F). We note that these published bulk cell Hi-C data were obtained from tissues that were first dissociated into cell or nuclei suspensions before performing the Hi-C steps, in contrast to our method with tissue sections. Hence, it may also be that the cell/nuclei isolation step might have contributed to the observed differences. Overall, thus, these results demonstrate that our tissue section Hi-C protocol is highly effective and may provide more faithful depictions of chromatin conformations in tissues.

### Histo-LCM-Hi-C for the characterization of defined cells in tissues

With our tissue section Hi-C procedure established, we next sought to isolate and characterize specifically defined cells using histological staining and LCM. To demonstrate the power of this method, we focused on characterizing the resident macrophages of the liver, the Kupffer cells, which comprise a significant minority of the total liver cells in mice (8 to 9%) [37]. As such, the chromatin conformation of these cells is expected to be not evident in data from bulk liver tissue samples. Indeed, we note that, to date, there is no data of the chromosome confirmation of these cells in the literature.

We first verified that the staining and LCM isolation did not significantly affect the Hi-C results compared with our tissue section results (SCC of 0.95) and yielded mapping data of comparable quality (Table S1). We then used LCM to select individual KC (**Figure 3A** and **3B**), using the F4/80 protein as a marker [38]. To be clear, owing to the randomness of the sectioning with regards to the positions of the cells and the similarity in size between a typical nucleus and the section thickness, we expected that each nuclear region identified in our sections was not an intact nucleus but only a portion of the nucleus. Indeed, we found that ∼600 of these nuclear regions yielded only about 1.63 ng DNA, which corresponds to about 270 intact cells. Experimentally, we found that we required ∼600 of these regions for the construction of an effective library for next-generation sequencing. In particular, examination of the DNA fragment size distribution revealed no or an abnormal distribution from the libraries obtained fewer than 600 nuclear regions (Figure S7A and 7B). Further, sequencing of this material at a relatively low depth to examine material quality (namely, total reads < 15 M) showed an extremely high proportion of duplicate reads (> 80%), revealing a very low library complexity. A simple calculation suggested that this lower cell limit appears to reflect a limitation of the Hi-C method, and not (for example) significantly inefficient enzymatic reactions under our conditions. Specifically, based on the total valid pairwise reads of one library obtained in our experiment (and assuming there were indeed effectively 270 cells), we found that our coverage per cell (103k valid pairwise reads) is comparable to that observed from typical single cell Hi-C studies (where there are clearly no access issues to the added enzymes) [39, 40] (Table S2). Thus, overall, we believe that our method exhibits a comparable level of efficiency to at least most single cell Hi-C assays.

**Figure 3.**
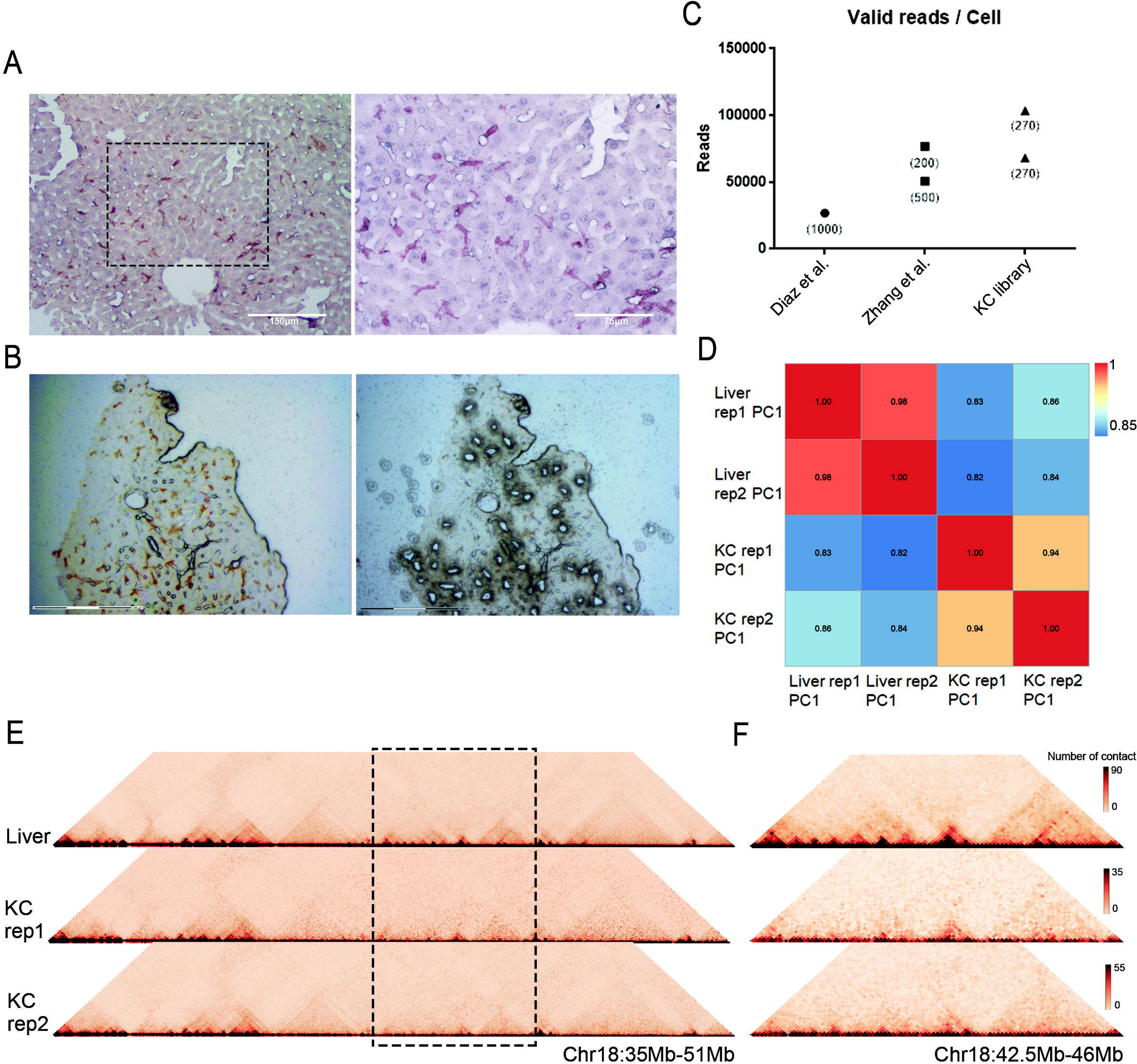
Generation of high quality Hi-C maps for specific rare cells in the liver. **A.** Immunohistochemistry images of KCs in the mouse liver section, using anti-F4/80 to label the KCs and Hematoxylin to label the nucleus. The right panel is a magnified image of the dotted boxed region in the left image. Scale bar: 150 μm (left) and 75 μm (right). **B.** A typical stained section before (left) and after (right) laser microdissection. Scale bar: 150 μm. **C.** The average valid pairwise reads per cell of the KCs Hi-C libraries compared with previously obtained low input Hi-C data [26, 41]. The numbers in the figure indicate the number of cells in each Hi-C library. **D.** Correlation results of the PC1 values from replicates derived from tissue section Hi-C libraries of the liver and KCs. **E.** Comparison of the contact maps generated from tissue section Hi-C of liver and KCs. **F.** A magnified region of the heatmap within the dotted boxed region of “E”. Resolution: 40 kb.

We obtained two libraries from a single mouse and found that the Hi-C maps from each sample were highly correlated (SCC of 0.96) (Figure S8A). To further validate the reproducibility of these results, we also obtained an additional four libraries from a different mouse. We found that these were also highly correlated with each other and with those of the first mouse (SCC of ∼0.93) (Figure S8A). Notably, the KC libraries that were obtained from as few as ∼300 cells exhibited higher valid interaction ratios compared to standard Hi-C protocols, underscoring the enhanced sensitivity of our method (Figure S8B). We find that under comparable sequencing depth, the data obtained using Histo-LCM-Hi-C were on par with other published low-input Hi-C methods, further supporting the idea that our Hi-C method is comparable to many of the established *in vitro* methods [26, 41] (**Figure 3C**; Table S3).

The data from each mouse were pooled together to obtain a KC replicate map (for each mouse). Examination of these maps at the level of compartments showed that the KC samples were much more similar to each other than to the total liver sample (**Figure 3D**). This was also evident by direct inspection of the Hi-C maps (**Figure 3E** and **3F**). Owing to the high consistency between the two biological KC replicates, to improve the resolution of the final map for more detailed analyses below, we combined the replicate data, ultimately obtaining a map at 40 kb resolution.

Notably, our method achieved this resolution with only ∼3000 nuclear regions (∼1500 intact cells), whereas standard Hi-C typically uses 10^6^ cells for comparable resolution. To provide insight into the relationship between cell number and resolvable structural features, we further examined the maps obtained from 300-cell, 750-cell, and 1500-cell samples. The resolution of these samples, following the conventional definition of resolution (80% of bins containing 1000 reads), is 200 kb, 80 kb, and 40 kb, respectively. We calculated the number of A/B compartments at a bin size of 200 kb for the 300-cell, 750-cell, and 1500-cell samples and found very similar numbers of compartments for each (Figure S9A). With the 750-cell and 1500-cell samples, we also calculated the number of A/B compartments at a bin size of 100 kb (since this is the typical bin size used for compartment identification) and found that there was a slightly greater number of compartments (Figure S9A), as expected when a smaller bin size is used. For the TADs, we compared the numbers from the 750-cell and the 1500-cell samples at a bin size of 80 kb and found ∼3500 TADs for both cell-sample sizes (Figure S9B). We also calculated the TAD number at a 40 kb bin size with the 1500-cell sample and found ∼5300 TADs. Thus, the majority of TADs can be resolved even with only 750-cells. Finally, for loops, we examined the 750-cell and 1500-cell sample at 80 kb bin size and the 1500-cell sample at 40 kb bin size using Mustache [40]. We found that only about half of the loops observed in the 1500-cell sample at 40 kb bin size were observed in the 750-cell sample at 80 kb bin size, clearly reflecting the improved signal-to-noise ratio and precision with the greater cell-number sample (Figure S9C). Thus, overall, these results suggest that with 300 cells, we can obtain reliable information about compartments, while 750 cells would be needed for reliable measurements of both compartments and TADs and 1500 cells would provide the most structural information about compartments, TADs and loops.

We also examined our 1500-cell KC map to evaluate the extent to which it resolved true chromatin interactions from background noise, compared to a larger-cell number sample (namely, liver). Using FitHiC2, we found that the data from both the liver and KC samples exhibit a similar level of such true contacts (Table S4) and both exceed other published work (Figure S10). We also examined our results after down-sampling the reads of small-cell-number (KC) datasets, but found no significant difference from the results without down-sampling (Table S4).

### Analysis of the KC chromatin architecture reveals mechanistic details underlying its functioning

KCs are the resident macrophages of the liver, and as such, they play crucial roles in immune reactions and liver-specific metabolic functions [42]. Their primary functions include clearing microorganisms and cellular debris from the blood, removing aging red blood cells, providing a barrier to prevent various pathogens and their toxic byproducts (such as endotoxins) from entering the systemic circulation, and supporting the protein and lipid metabolism [43].

Overall, our final KC Hi-C map exhibited many features that are similar to those that have been observed in previous *in vitro* Hi-C studies of other cell types [44, 45], such as, for example, showing ∼5000 TADs that exhibit a median size of 400 kb (Table S5). We were particularly interested in examining those regions of the maps that are different between the KCs and the whole liver, as these might reveal novel details of gene regulation within the KCs. Owing to the established relationship between compartment type and gene expression, we first compared these maps at the compartment level (determined at a bin size of 100 kb), finding 5% (1270) of regions that are A in KCs and B in the whole liver and 6% (1671) that are B in the KCs and A in the liver (Figure S11A). We also note that for those regions that were unchanged in compartment type, we found a difference in overall number of contacts within each compartment, which thus indicates a significant different level of compaction within each compartment between these samples (Figure S11B).

We found that the regions that are A in the KCs and B in the liver are enriched for immune-related genes. In particular, we found an enrichment of genes involved in pathways such as the activation of innate immune response and positive regulation of lymphocyte proliferation (**Figure 4A**, Table S6). For example, we found that both *Ccl2* and *Il16* are located in the A compartment in the KCs and B in the liver; both of these proteins are pro-inflammatory cytokines that are chemotactic for monocytes, CD4+ T lymphocytes and eosinophils [46, 47]. Thus, these results suggest that the expression of genes of this important pathway of KCs for hepatic immunity is indeed regulated by specific chromatin conformations. Likewise, we note that those regions that are B in the KCs and A in the liver were more often associated with genes involved in core metabolic functions of the liver, which are not expressed in the KCs, also likely related to their location within the B compartments in the KCs (Figure S11C).

**Figure 4.**
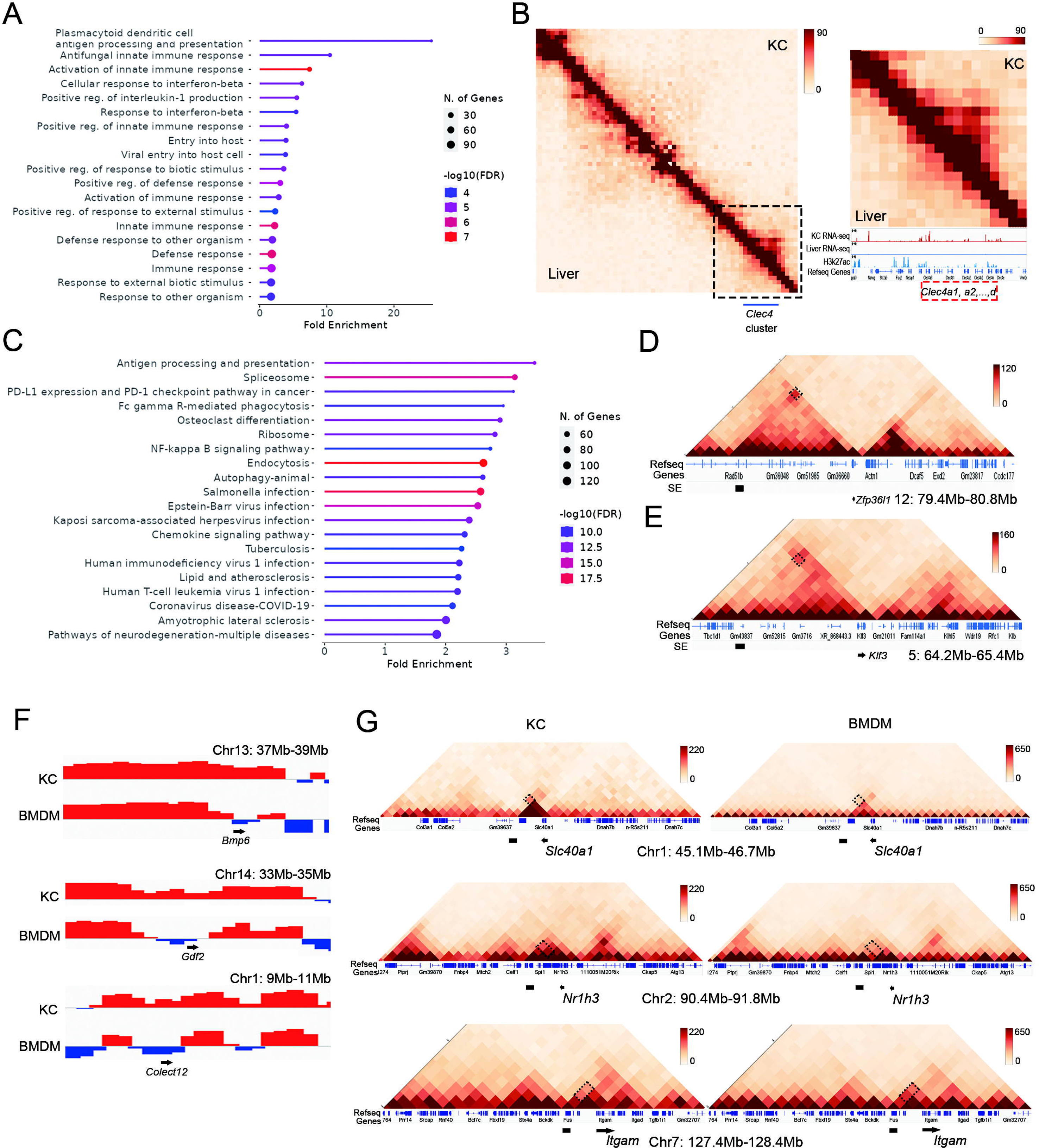
KC specific chromatin structures and comparison with BMDM. **A.** GO analysis results of the genes in the KC A compartment and liver B compartment. **B.** Visualization of contact maps generated from tissue section Hi-C of the KCs (top) and liver samples (bottom) in Chr6, from 120.9 Mb to 123.4 Mb, resolution: 40 kb. The right panel is a magnified heatmap of the boxed region in the left panel from 122.6 Mb to 123.4Mb, together with RNA-seq of KCs and liver tissue, H3K27ac ChIP-seq data of the KCs and Refseq Genes. **C.** GO analysis results of the genes having strong contacts (from FitHiC2, q-value < 0.005) with SEs in the KCs. **D.** and **E.** Visualization of contact maps showing the SE-gene contacts with *Zfp36l1* (**D**) and *Klf3* (**E**), resolution: 40 kb. In both images, the black bar indicates the position of the SE, together with Refseq and SE. SE, super enhancer. **F.** Plots of the PC1 values reflecting compartment-level interactions from the KCs (upper panels) and BMDMs (lower panels). Positive values (red) reflect A compartment and negative values (blue) reflect the B compartment. **G.** Visualization of the contact maps of three exemplary regions, showing stronger contacts between putative enhancers [53] (the black bar) and genes with prominent functions in either KCs or BMDMs, together with the Refseq genes. Resolution: 40kb.

We performed diffHic [21] to identify regions of local differences in structure between the KC and total liver samples. Overall, we found that ∼84% of the genome (at 40 kb bins) exhibited at least one change in contact with another region of the genome (Table S7). One of the most significantly different loci overlaps the *Clec4* gene cluster, whose members are specifically expressed in KCs [48, 49] and involved in phagocytosis and production of inflammatory cytokines and chemokines in KC [50–52]. We find that this locus forms a separate TAD in the KCs that is completely absent in the total liver sample (**Figure 4B**), which may ensure effective contacts between enhancers and the *Clec4* promoters in the KCs. Another top diffHiC loci overlaps the *Ifi200* family gene cluster that exhibits significantly greater contacts with a putative enhancer [53] in the KCs than in the liver (Figure S12A and S12B). This gene cluster encodes PYHIN proteins, which play important roles in sensing pathogen DNA [54], cytokine promoter induction in macrophages [55] and cell proliferation [56], all prominent functions of KCs in the liver.

One of the most useful details provided by Hi-C is the identification of statistically significant contacts between enhancers and gene promoters that underlies the promotion of expression by the enhancers [57, 58]. Without such data, it is generally assumed that an enhancer regulates the closest expressed gene within a specific genomic size, usually 200 to 250 kb. While our dataset provides this resource for all putative enhancers, we were particularly interested in identifying the contacts between super-enhancers (SEs) with the gene promoters in our KC Hi-C map. SEs are thought to ensure a high level of expression of specific genes that play a significant role in functions related to the phenotype of a cell [22]. Thus, we expected that the identification of these contacts in our data would reveal the means of regulation of a core group of critical KC genes. We used published H3K27ac ChIP-seq data from KCs [59] to identify the SEs of KCs, and used FitHiC2 to identify the most significant contacts between the KC SEs and the transcription start sites of expressed genes in the KCs (Table S8). We found that the median distance between these TSSs and the SEs that they contact is 200 kb (Table S8). Hence, at least half of these contacts would likely be missed without this direct evidence of contact between these loci. We found that, confirming our expectations, these genes were predominantly enriched for immune-related signaling pathways, including antigen processing and presentation and the NF-kappa B signaling pathway (**Figure 4C**). For instance, the *Zfp36l1* gene shows a significant interaction with an SE that is approximately 500 kb away (**Figure 4D**), well beyond the 200 kb to 250 kb conventionally expected range for enhancer/promoter contacts. Similarly, the *Klf3* gene interacts significantly with a SE located 400 kb away (**Figure 4E**). Both of the proteins play significant roles in KC functioning, with Zfp36l1 involved in cellular metabolism [60] and Klf3 playing a dominant role in NF-κB-mediated inflammation [61]. Notably, the interaction of the former with the SE is not prominent in the liver Hi-C data (Figure S12C and S12D).

Along the same lines, previous work that examined differential expression among several different tissue-resident macrophages identified a signature KC-specific gene set [62]. We found that nearly half of these KC-specific genes (144 out of 303) made specific contacts with SEs in KCs (Table S9). Among these is Clec4f, one of the marker-defining genes of the KCs [63, 64]. We find that this gene exhibits long range contacts with 2 SEs that are 300 to 600 kb away (Table S8). We note that earlier work identified a putative enhancer for the *Clec4f* gene in KCs that is approximately 20 kb from the TSS of this gene [65]. Our data suggest a more complicated means of regulation of this gene that involves multiple distant SEs. The strict expression of this gene to KCs [65] (absent in other liver cells or extrahepatic macrophages) is thus likely owing to these chromatin interactions with the distant SEs. In addition, we also find that one of the most highly expressed groups of KC-specific genes, the members of complement C1q proteins (C1qa, C1qb, and C1qc) exhibits many distal contacts with an SE located around 450 kb away (Figure S13). A putative enhancer for these genes in a number of human immune cells has been identified [66], but this enhancer location does not appear to be syntenic with the SE that we find contacts these genes in mouse KCs. This suggests that there is a KC-specific means of regulating the *C1q* genes by specific chromatin structures that is important for systemic immune surveillance and pathogen elimination [67]. Thus, overall, our maps indeed enable identification of a core group of KC genes whose expression is likely owing to contacts with SEs identified in these Hi-C maps that can now be tested with additional functional assays.

We also note that most KCs are located within zone 2 (between the periportal and pericentral regions) [68] and there is some suggestion that, like the hepatocytes, the KCs may also exhibit zonated expression. Thus, future work aimed at characterizing the chromatin structure spatial dependence in the liver, together with transcriptomic data obtained from similar cells, could provide significant insight into the role of local microenvironment on these cells and their influence on liver function.

Finally, we expected that our data might also be useful to illuminate differences in regulatory mechanisms among different types of macrophages [69, 70]. While to date, to our knowledge, there are no Hi-C profiles of other tissue-resident macrophages that were directly isolated from native tissue, there is a profile of bone marrow-derived macrophages (BMDMs) that were induced from BM derived monocytes *in vitro* [71]. This model system is often considered to exhibit many of the characteristics of macrophages in the blood [72, 73]. Hence, we expected that a comparison between our KC Hi-C data with this published data might provide insight into liver-specific regulatory mechanisms for macrophages

Interestingly, these maps exhibit a similar level of overall correlation (SCC of 0.82) as that between the KCs and our total liver sample (0.86), thus indicating a substantial level of differences in structure between these two types of macrophages. In total, 12% of compartment bins and 42% of TAD borders (Figure S14A) were different between the two cell types. Genes in the A compartment in the KCs and B compartment in the BMDMs are enriched for pathways related to monoatomic cation (including iron) homeostasis and intracellular iron ion sequestration (Figure S14B), both of which are functions of KC involved in recycling aged erythrocytes [74–76]. For example, *Bmp6* and *Gdf2*, play a significant role in maintaining iron homeostasis and are all in the A compartment in the KCs and the B compartment in the BMDMs (**Figure 4F**). Further, the prominent scavenger receptors, *Colect12* (Figure 4F) and *Marco* (Table S10), are also in the A compartment in KCs and the compartment B in BMDMs. Recent work has shown that scavenger receptors genes contribute to a higher accumulation of low-density lipoproteins in KCs, a unique function of the liver-resident KCs [63, 77].

We identified significantly different regions of contact between the KCs and BMDMs using diffHic [21] and identified several instances that appear to underlie the differences in gene expression (Table S11). For instance, the *Slc40a1* gene, which encodes for ferroportin that plays a pivotal role in iron metabolism in KCs [75, 78], makes specific contact with a distal putative enhancer [53] only in the KC Hi-C map (**Figure 4G**). Additionally, the *Nr1h3* gene, which encodes the transcription factor LXRα [62], also exhibits distinct interactions with an enhancer (in fact, an SE) in the KC map (Figure 4G), a chromatin architectural feature that likely drives its spatiotemporal expression. LXRα has been suggested to a lineage-determining transcription factor (LDTF) of the KCs [79], and we indeed find that other LDTFs (*Spic*, *Id1*, *Id3*, and *Irf7* [65, 80]) all also make significant interactions with distant SEs in the KCs (Figure S15, Table S8). For example, *Id3* is essential not only for KC differentiation during embryonic development but also for their anti-tumor activity in adulthood [81]. Therefore, the mapping of such chromatin structural architectures (as achieved here) provides novel insight into the mechanistic basis underlying expression and maintenance of the KC phenotype.

However, we also note that this comparison also enables identification of unique contacts only in the BMDM Hi-C data. For example, the *Itgam* gene that encodes the BMDM-specific marker Cd11b [82] exhibits a much stronger contact with a putative enhancer [53] in the BMDM Hi-C data (Figure 4G). Thus, beyond informing regulatory mechanisms within the KCs in particular, our data can provide insight into potential mechanisms of regulation within macrophages that function in other regions of the body, once similar data has been obtained from these regions.

## Discussion

The recent success of spatial transcriptomics methods in revealing previously unknown details of cellular functioning in tissues has driven the significant interest in the development of techniques that can provide complimentary data at the cellular resolution within native tissues [83]. Among such data, chromatin structure is now well known to play a significant role in the regulation of gene expression. Indeed, there are now a number of methods that can reveal regions of chromatin accessibility (notably ATAC-seq) directly from cells within tissues [84, 85]. As increased chromatin accessibility is associated with active enhancers and promoters, these studies have provided molecular-level insight into the precise transcription factors that underlie the gene expression in tissues [84, 86]. However, it is often not clear precisely which enhancer(s) regulate which gene(s) from this data. For this, identification of chromatin contacts, using methods such as Hi-C, would appear to be indispensable [87–89].

In this work, we provide the first Hi-C-based method to characterize the chromatin structure within specifically defined cells in the tissue. We show that, in terms of the Hi-C procedure, our method yields a data quality comparable to what is obtained *in vitro* (Table 1) and with a similar level of efficiency as published single cell Hi-C data (Table S2). As both LCM and Hi-C are common methods in many labs, we anticipate that our method will be rapidly adopted to provide direct information about the important chromatin contacts that underlie the functioning of cells within many different contexts. For example, this method could be used to detect specific cells within the tumor microenvironment, such as immune cells in close proximity to malignant cells exhibiting unique phenotypes, to enable not only a more comprehensive understanding of these phenotypes but also their potential cell-cell interactions. Similarly, Histo-LCM-Hi-C could be used to resolve dynamic chromatin reorganization during organogenesis. For instance, isolating SOX2+ progenitor cells [90, 91] at precise developmental stages within the subventricular zone of embryonic cerebral cortex could reveal how chromatin conformation contributes to the regulation of neurogenesis during mouse organogenesis.

We anticipate that there will be considerable interest to integrate Hi-C with other extant spatial -omics methods to provide a more comprehensive understanding of tissue functioning. In particular, our Hi-C method achieves full genome structure characterization of phenotypically-defined cell types. Spatial ATAC-seq approaches [90, 92] provide highly complementary information: they typically identify open chromatin loci (∼1 kb in breadth) that can be used to identify transcription factor binding sites within highly localized regions within the tissue. Spatial transcriptomics can map gene expression of at least 10,000 genes within the spatial context of tissues [2, 93], and spatial proteomics can be used to determine the protein composition of ∼tens of proteins per cell type within a tissue [94]. Still, spatial ATAC-seq and most spatial transcriptomic approaches are somewhat limited in their spatial precision, identifying features within spots that may contain more than 1 cell. In addition, spatial proteomics is limited by the number of proteins that can be simultaneously detected in each cell. By contrast, a major limitation of LCM-based methods is throughput. Thus, we believe that the integration of the greater-depth but lower-breadth of LCM-based methods with the existing lower-depth but greater-breadth capabilities of extant spatial -omics methods will prove to be much more effective in resolving basic features of tissue function than any method alone.

Yet a particular noteworthy aspect of our method, even among spatial -omics methods, is the ability to characterize cells that are a significant minority of the total tissue cell population. This is a simple consequence of the basic LCM method of identifying and isolating specifically defined cells in the tissue. We believe that the ability to characterize such rare cells within the tissue will indeed prove to be an attractive feature that will be exploited to compliment other high throughput based spatial -omics methods. We have previously established a method to characterize the transcriptome from specifically defined cells with LCM from as few as 50 cells [95] and other researchers have characterized other molecular properties, including metabolites and proteins, using LCM from tens to hundreds of cells [96, 97]. Thus, we envision that LCM-based methods will find a particularly attractive niche among the repertoire of spatial -omics methods by enabling a more precise understanding of specific cell populations that will be suggested to be of interest from other spatial approaches. Ultimately, such a level of understanding may be needed for all of the cells in the tissue to fully understand tissue behavior. However, until high-resolution, spatial, multi-modal methods are developed, the combination of the high-resolution data from a few specific cells using LCM-based approaches combined with lower-resolution higher throughput methods may prove to be a powerful approach that can now be employed with present technology.

## Ethical statement

The experiments with mice were approved by the Institutional Animal Care and Use Committee (IACUC) at Shanghai Jiao Tong University and approval number is A2020109.

## Data availability

The raw sequence data reported in this paper can be accessed at https://dataview.ncbi.nlm.nih.gov/object/PRJNA1187272?reviewer=u0joc2qr6rijib7ll50ro8id9m, and in the Genome Sequence Archive [98] in National Genomics Data Center [99], China National Center for Bioinformation / Beijing Institute of Genomics, Chinese Academy of Sciences (GSA: CRA024256) that are publicly accessible at https://ngdc.cncb.ac.cn/gsa.

## CRediT author statement

**Yixin Liu**: Investigation, Methodology, Writing - original draft. **Min Chen**: Software, Formal analysis, Writing - review & editing. **Xin Liu**: Investigation, Methodology. **Zeqian Xu**: Formal analysis. **Xinhui Li**: Investigation. Yan Guo: Investigation. **Daniel M. Czajkowsky**: Conceptualization, Formal analysis, Writing - original draft. **Zhifeng Shao**: Conceptualization, Methodology, Writing - original draft. All authors read and approved the final manuscript.

## Acknowledgments

This work was supported by grants from the National Key R&D Program of China (No. 2020YFA0908100), the National Natural Science Foundation of China (Nos. 31971151, 81627801, 81972909, and 32370572), and the K.C. Wong Education Foundation (H.K.). We thank Ming Cheng for his help with these experiments.

