## Supplementary figures and images for "Histo-LCM-Hi-C reveals the 3D chromatin conformation of spatially localized rare cells in tissues at high resolution"

### Supplemental figure 1

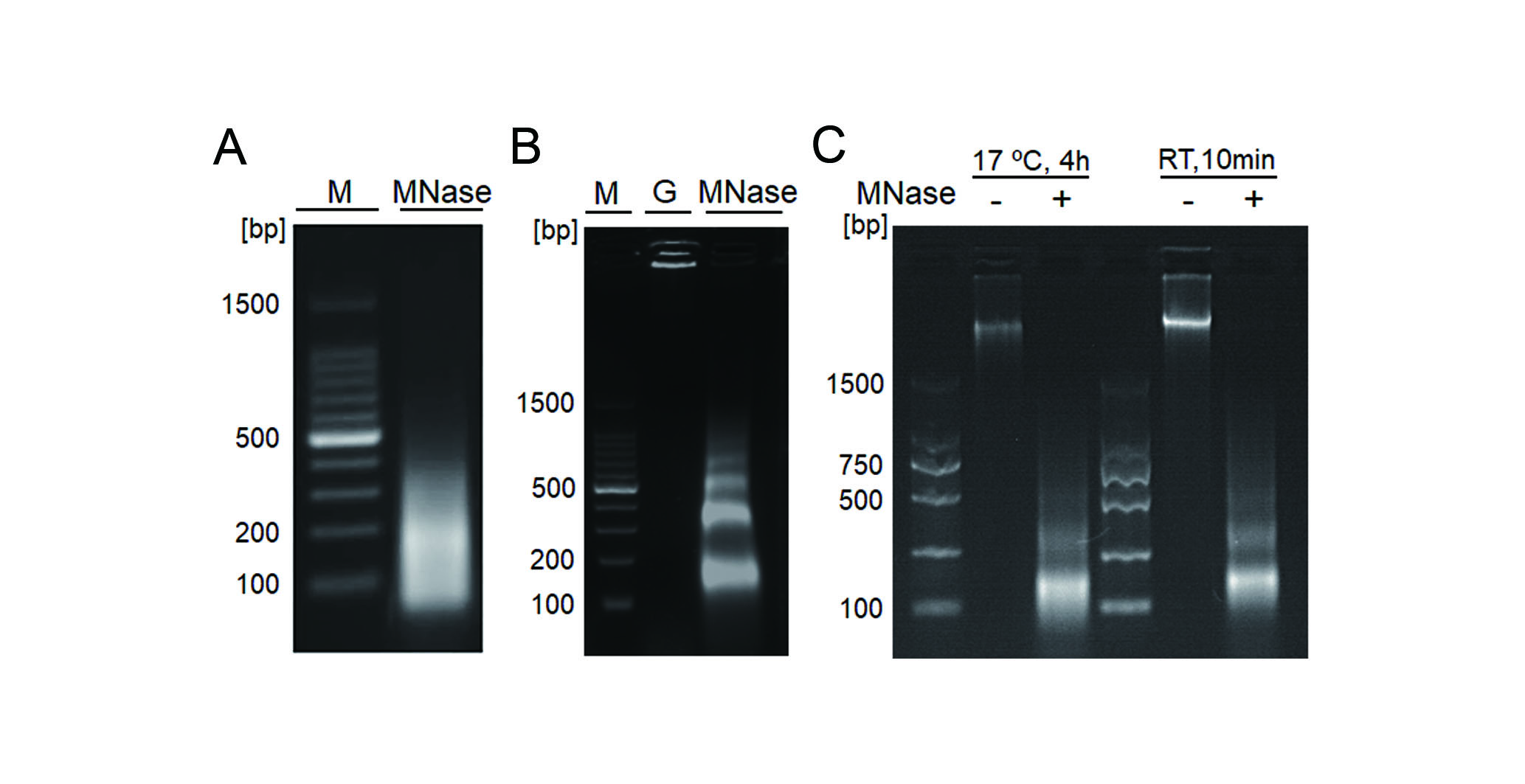

### Supplemental figure 2

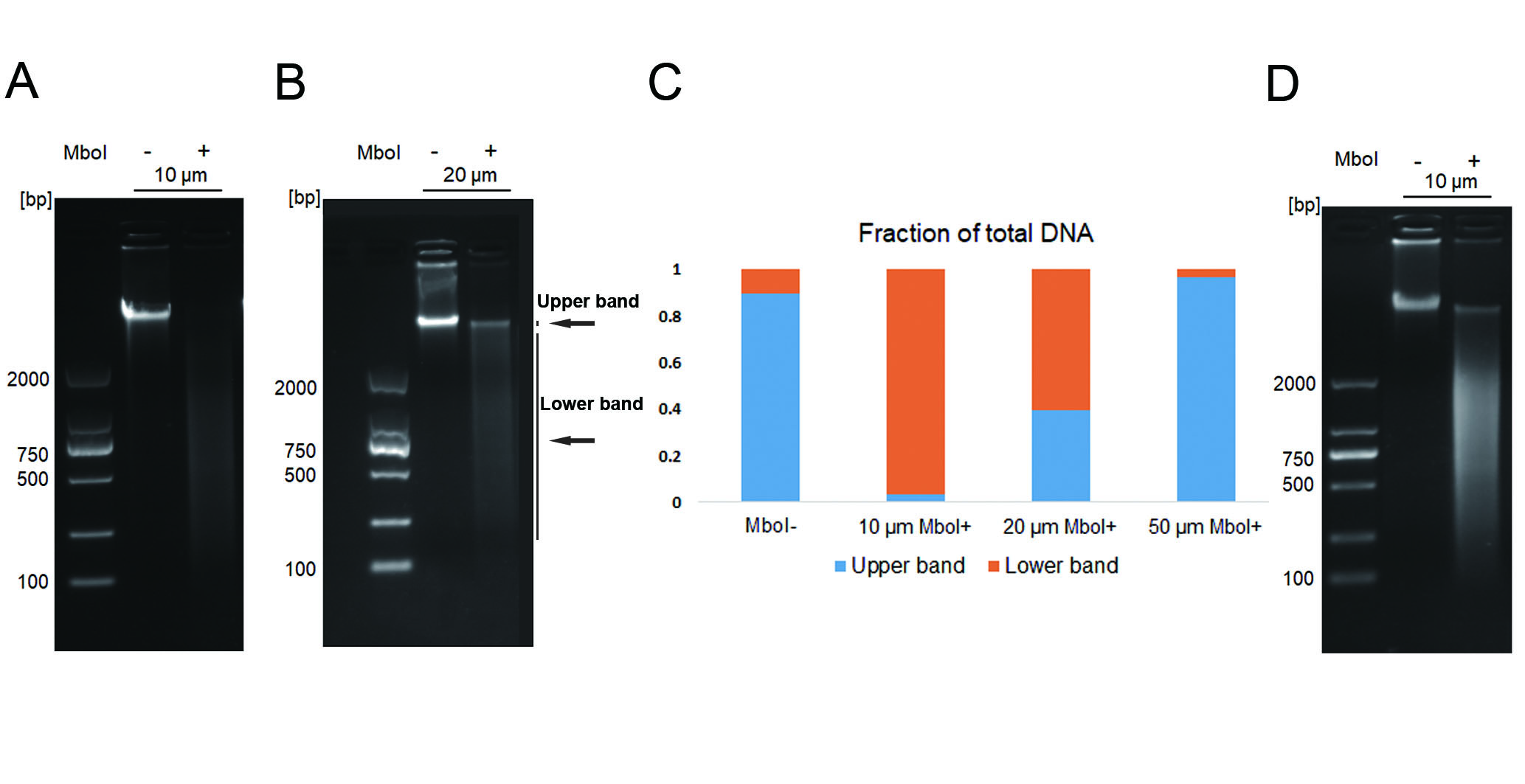

### Supplemental figure 3

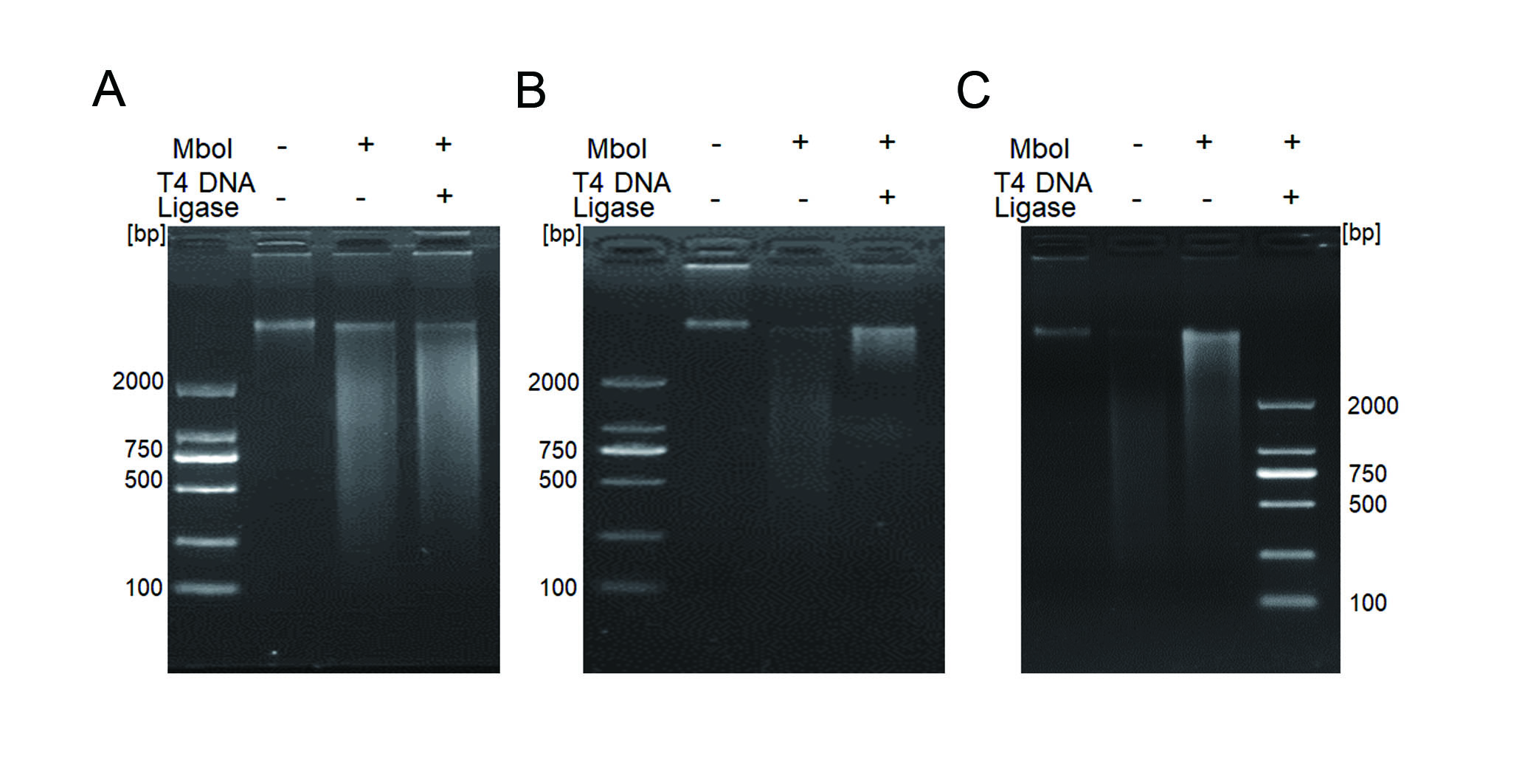

### Supplemental figure 4

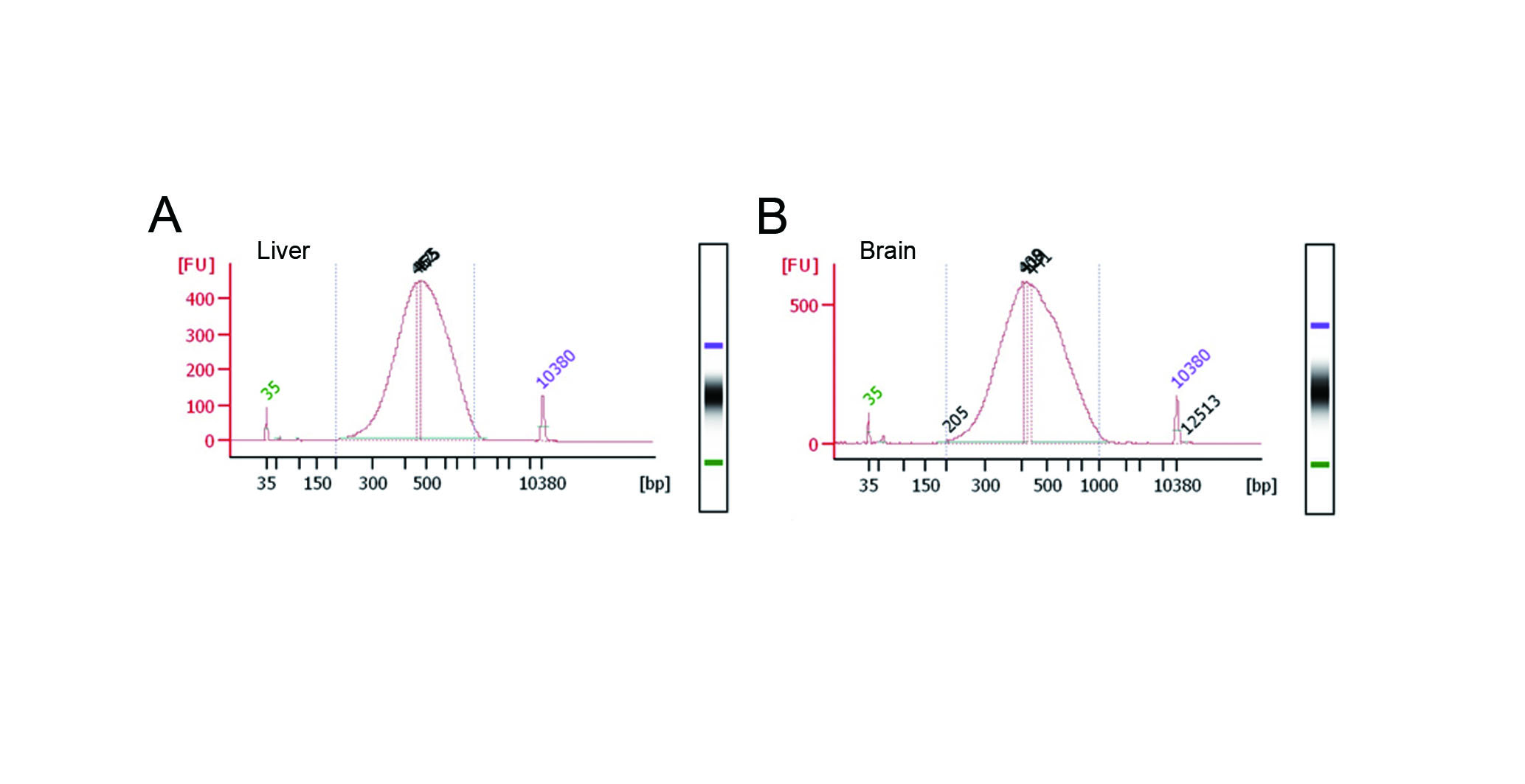

### Supplemental figure 5

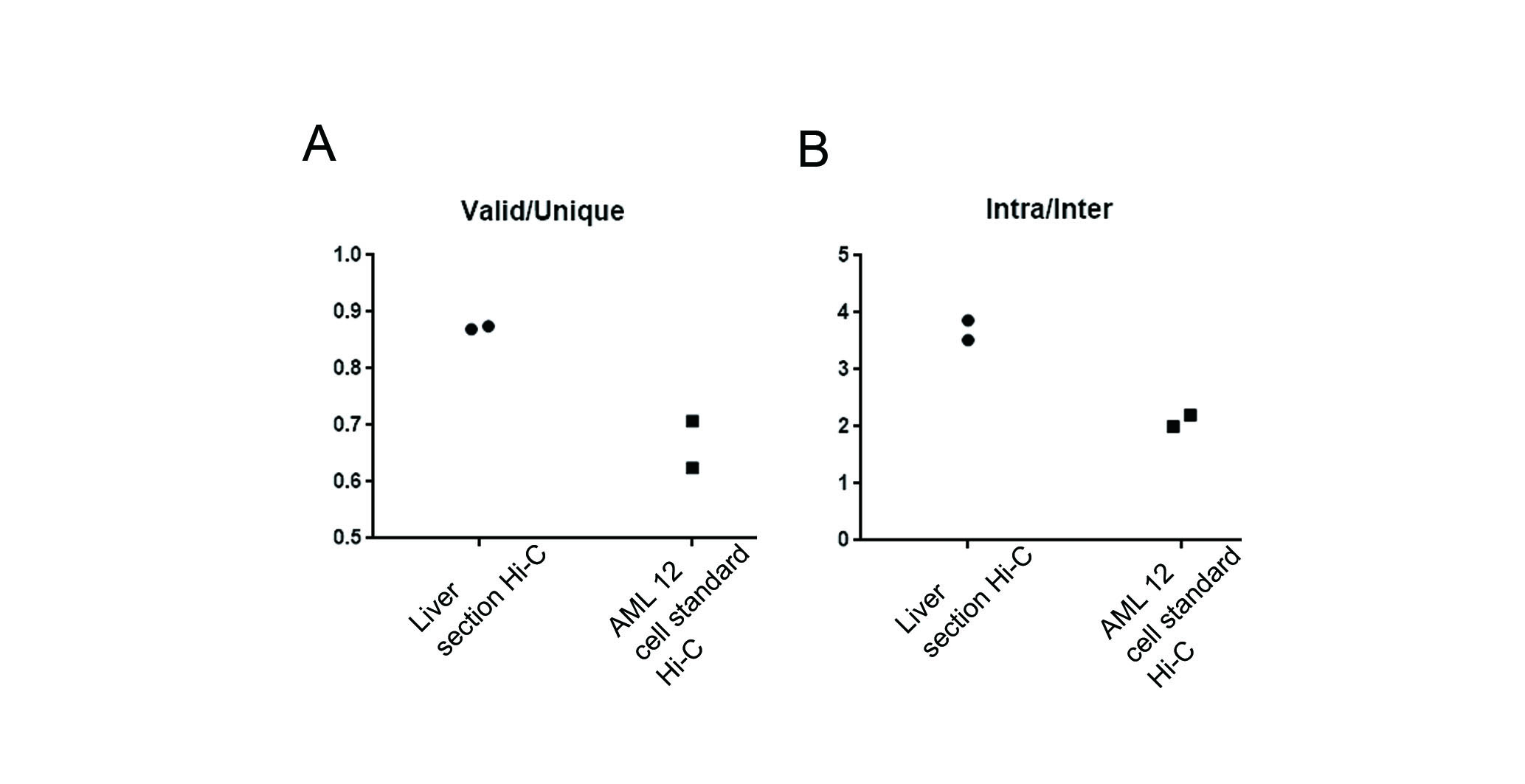

### Supplemental figure 6

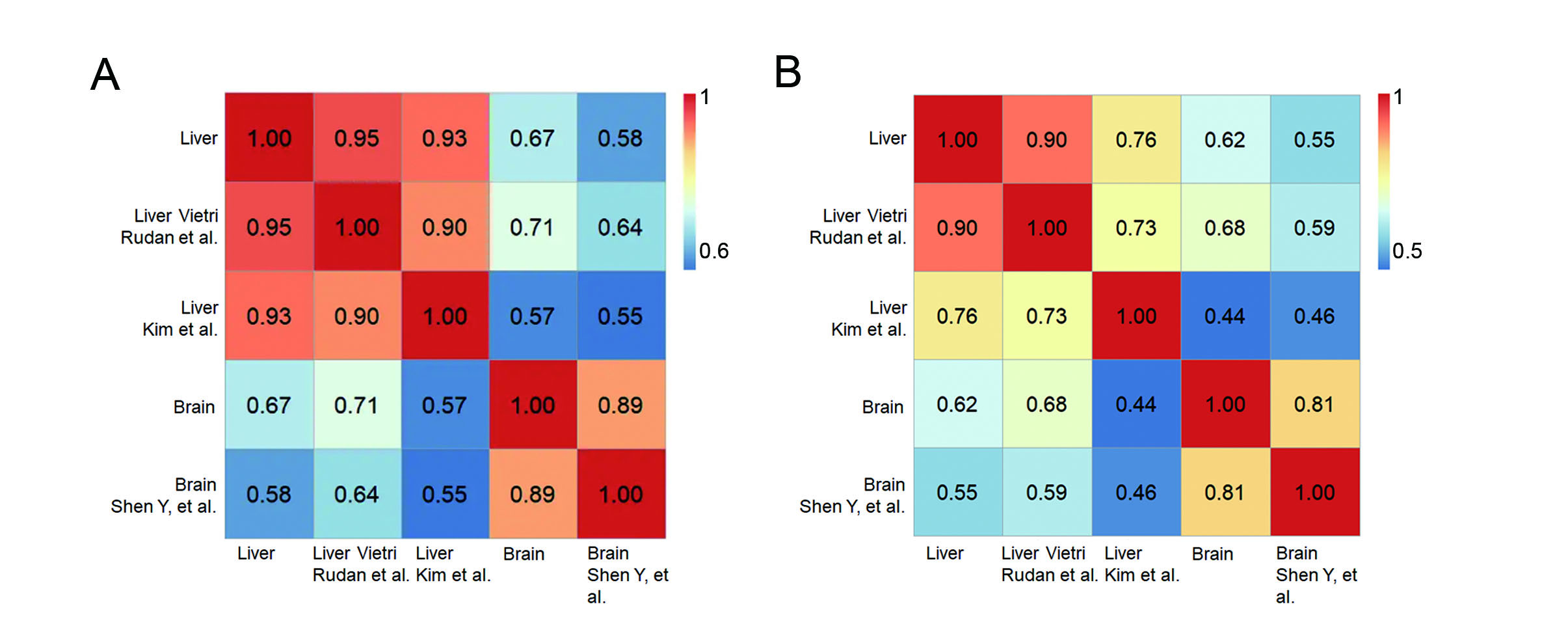

### Supplemental figure 7

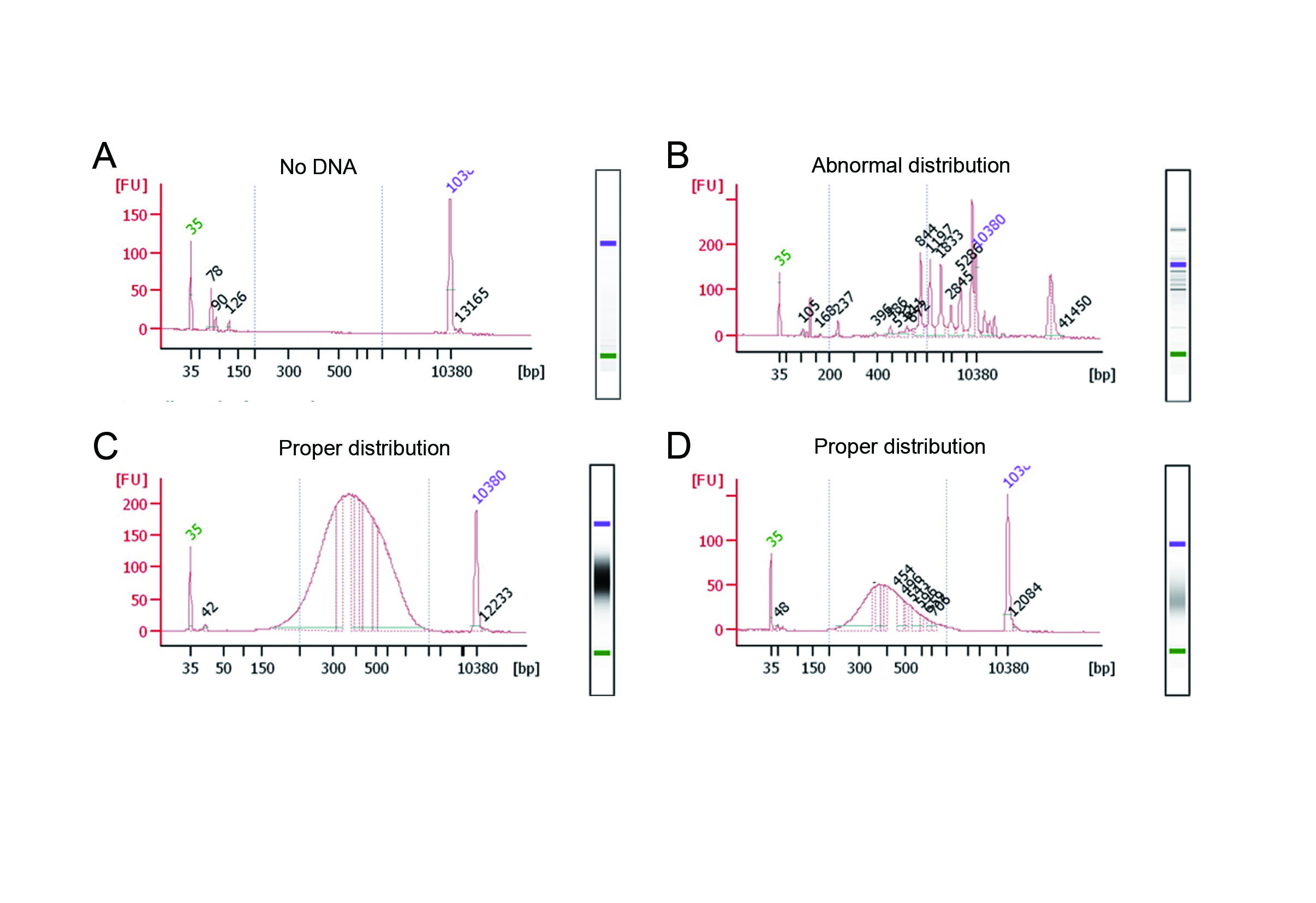

### Supplemental figure 8

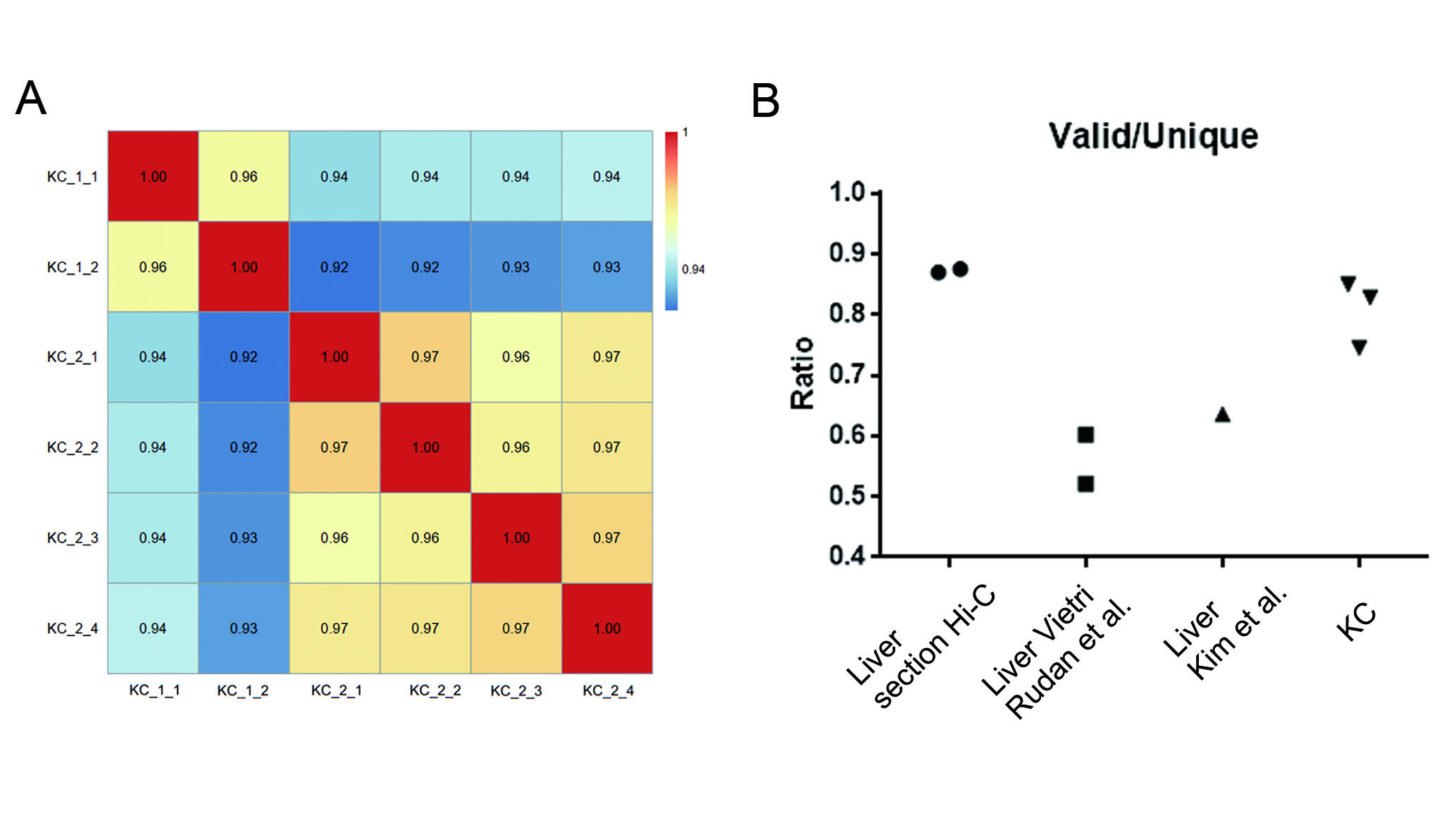

### Supplemental figure 9

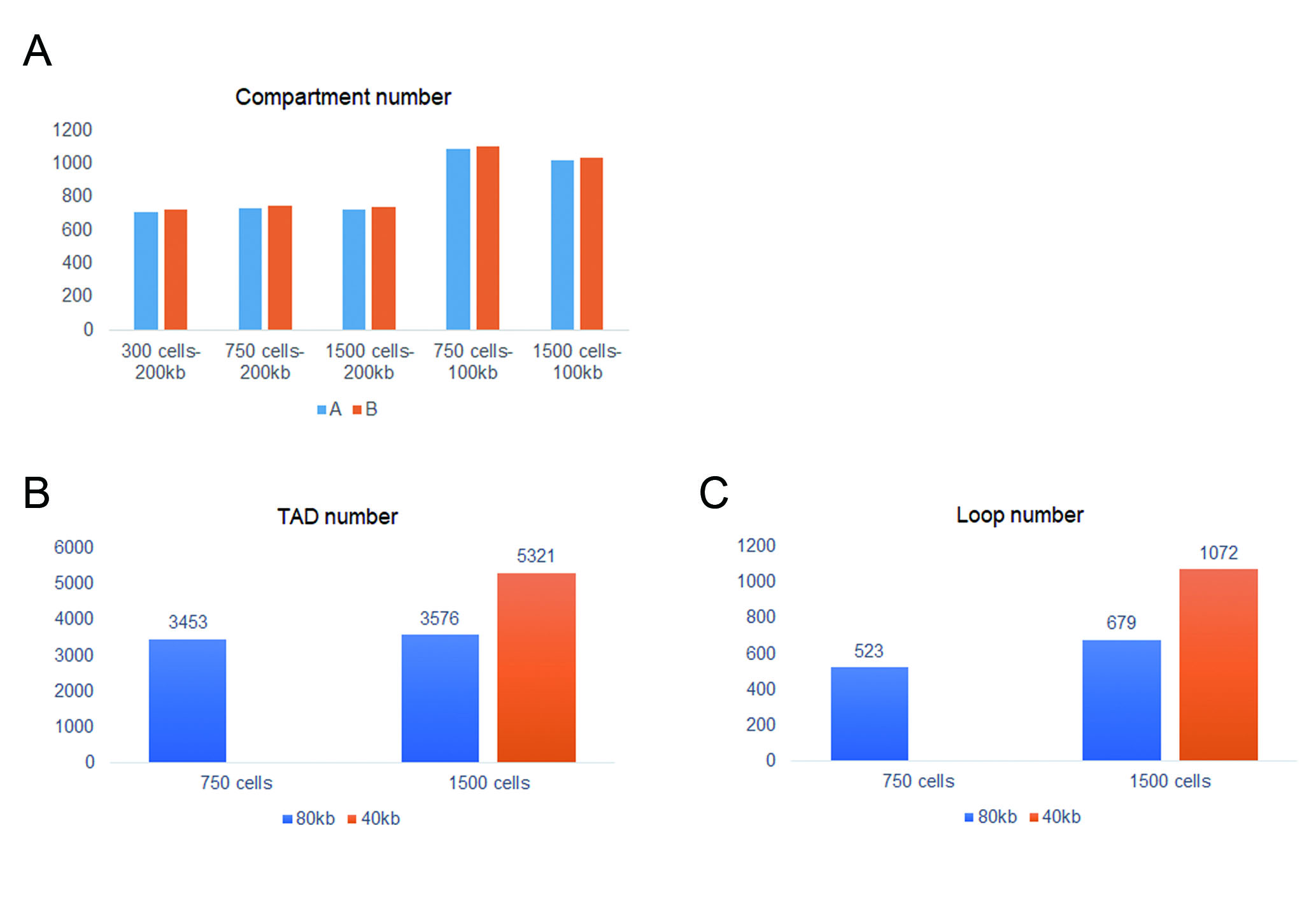

### Supplemental figure 10

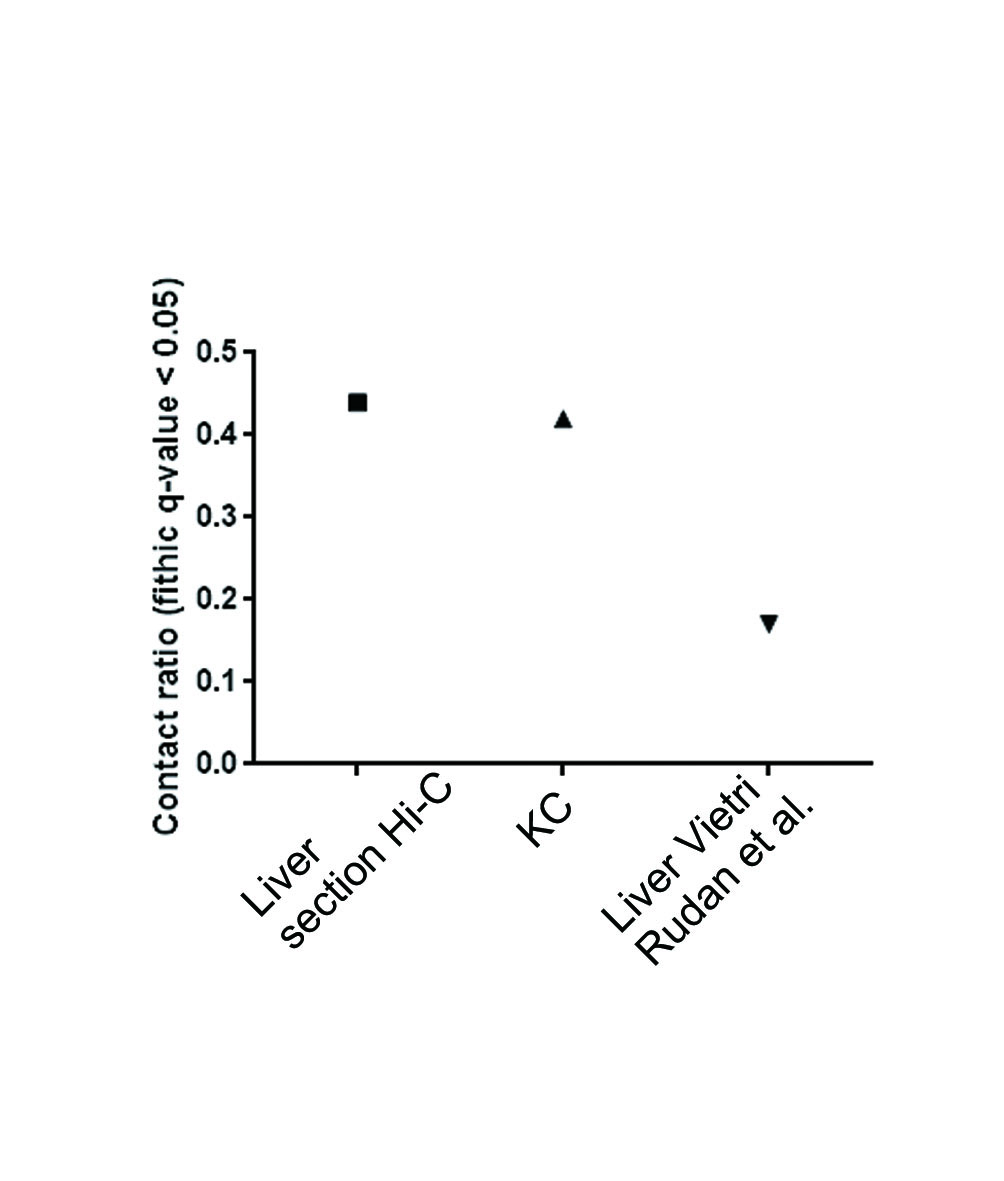

### Supplemental figure 11

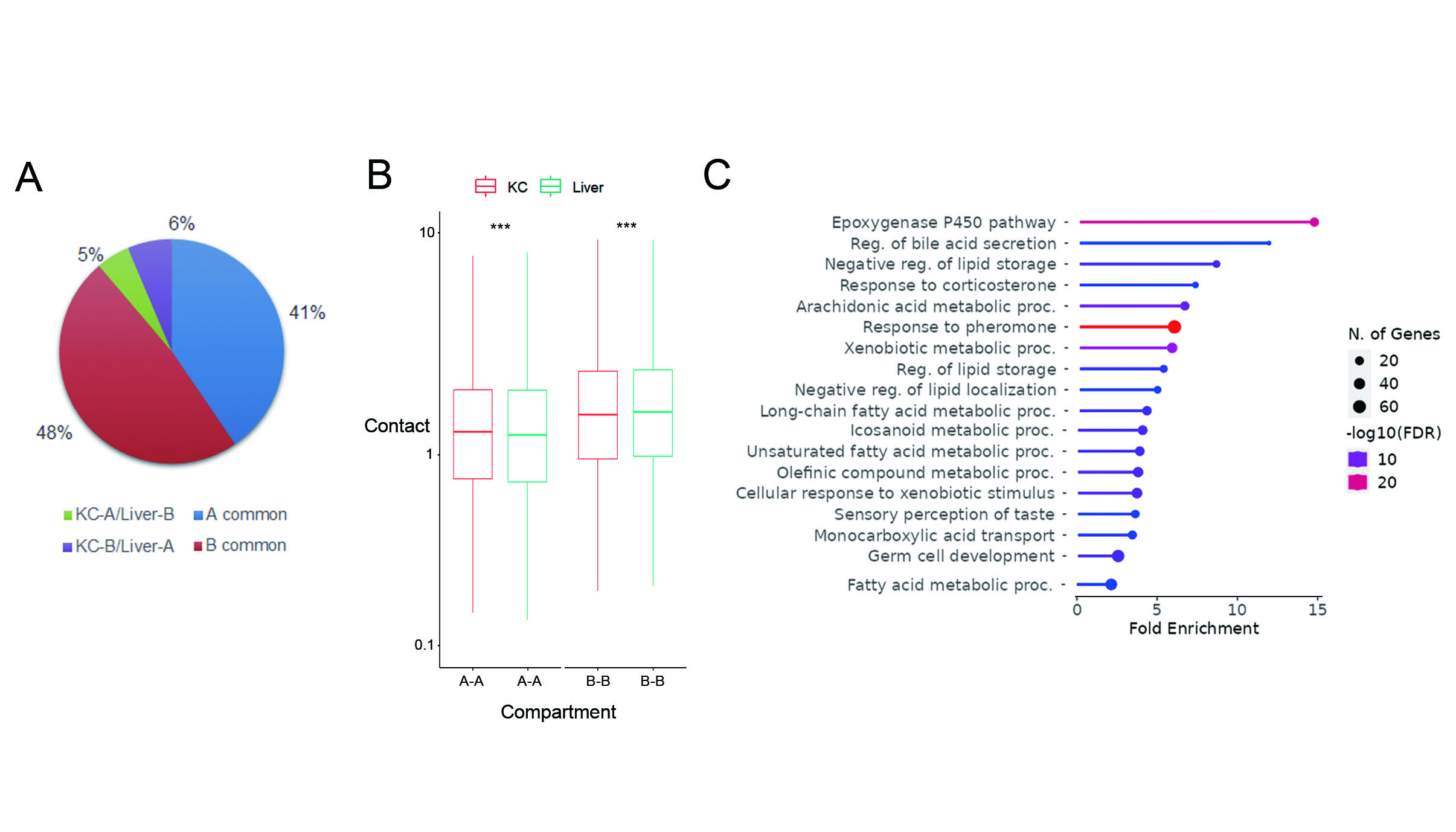

### Supplemental figure 12

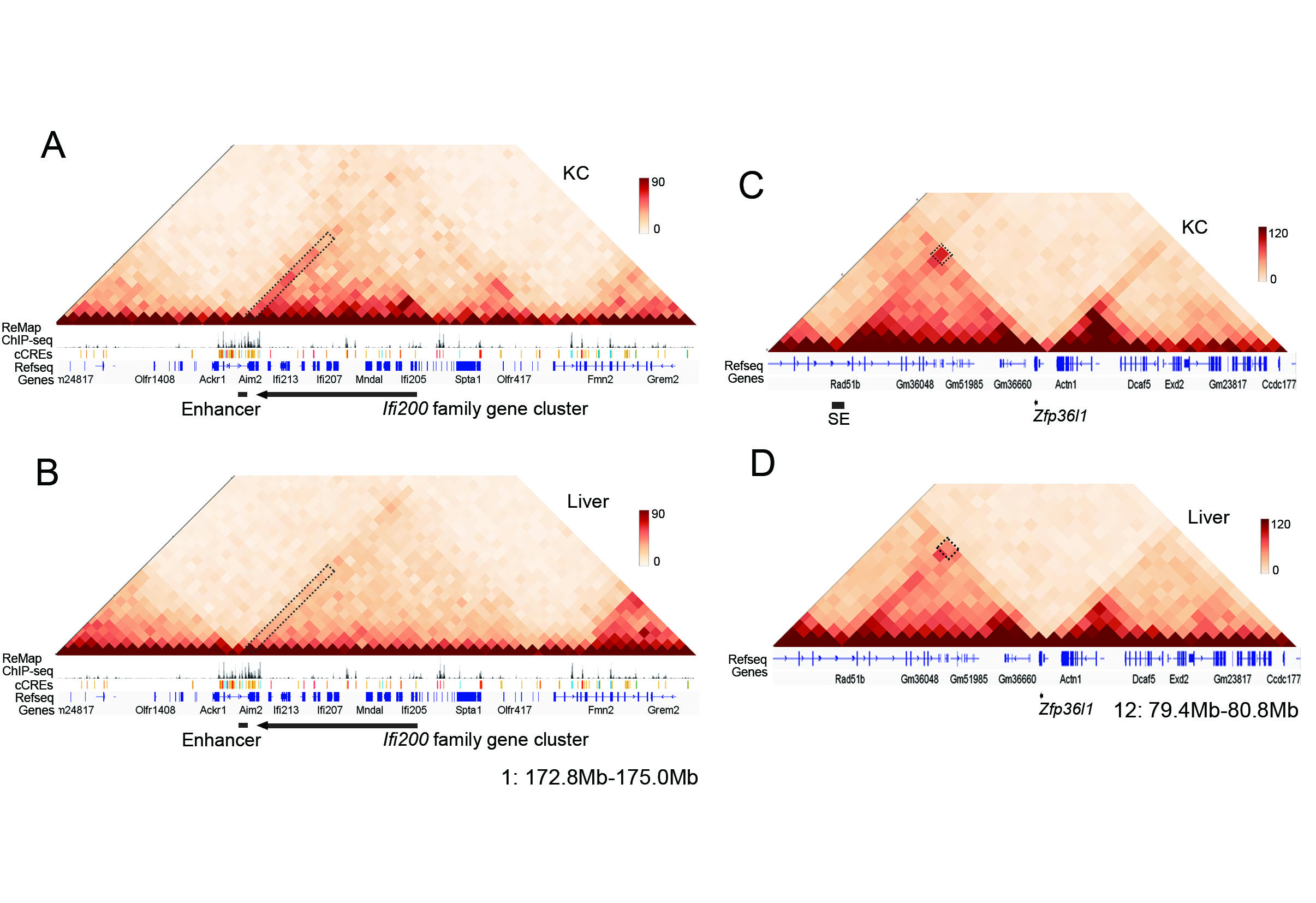

### Supplemental figure 13

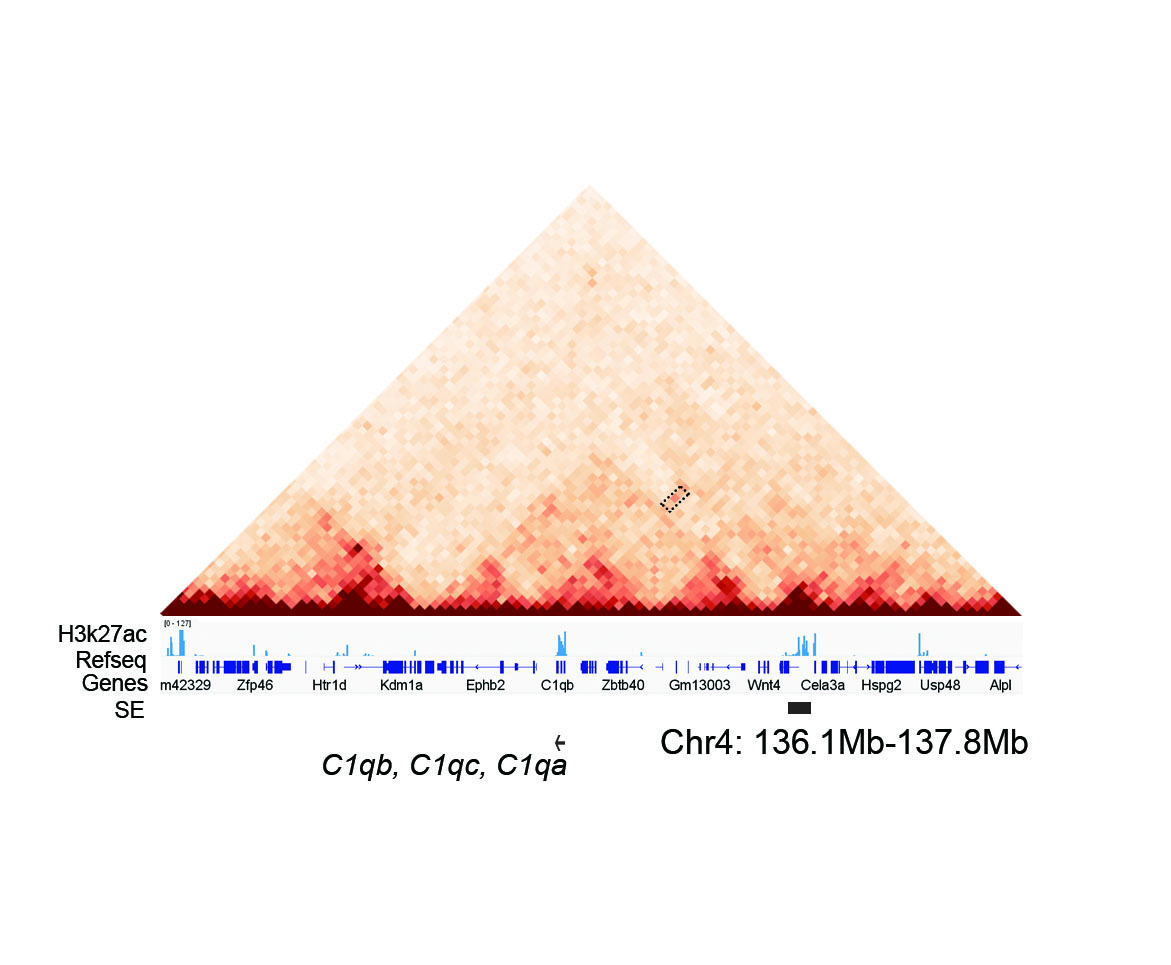

### Supplemental figure 14

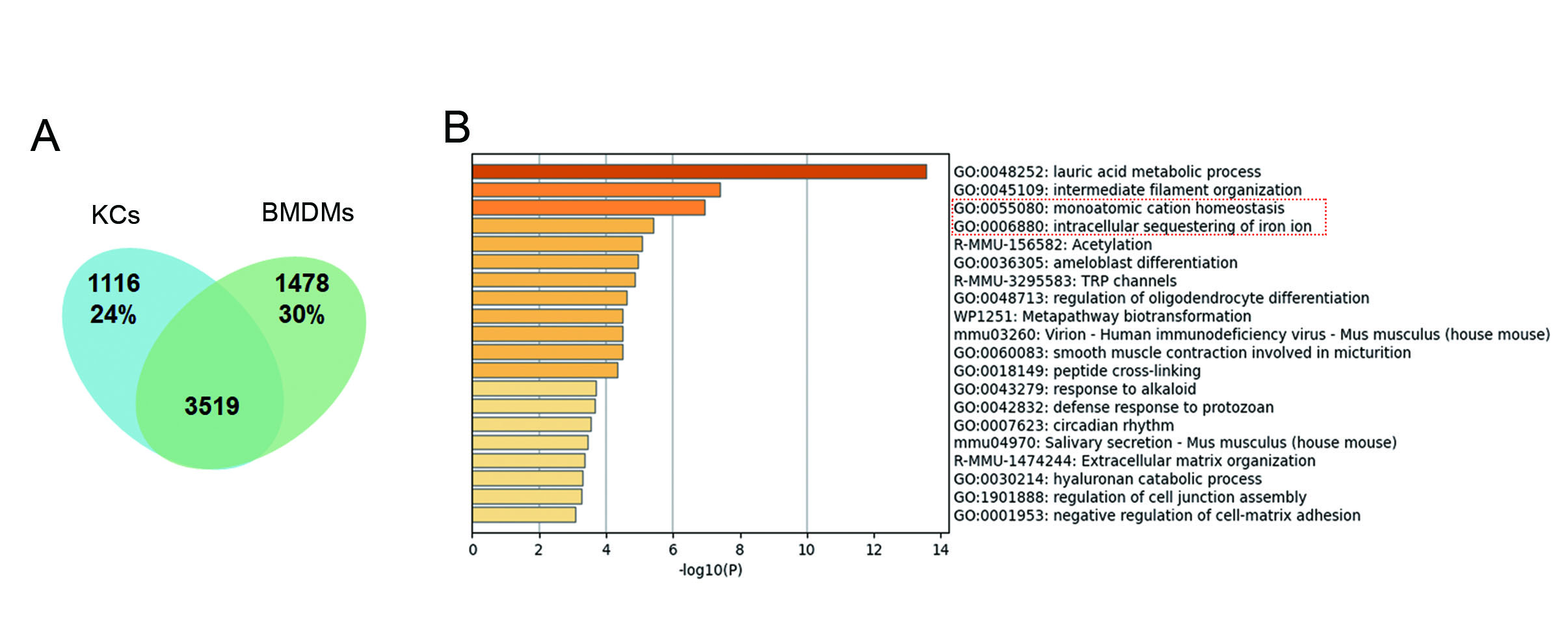

### Supplemental figure 15

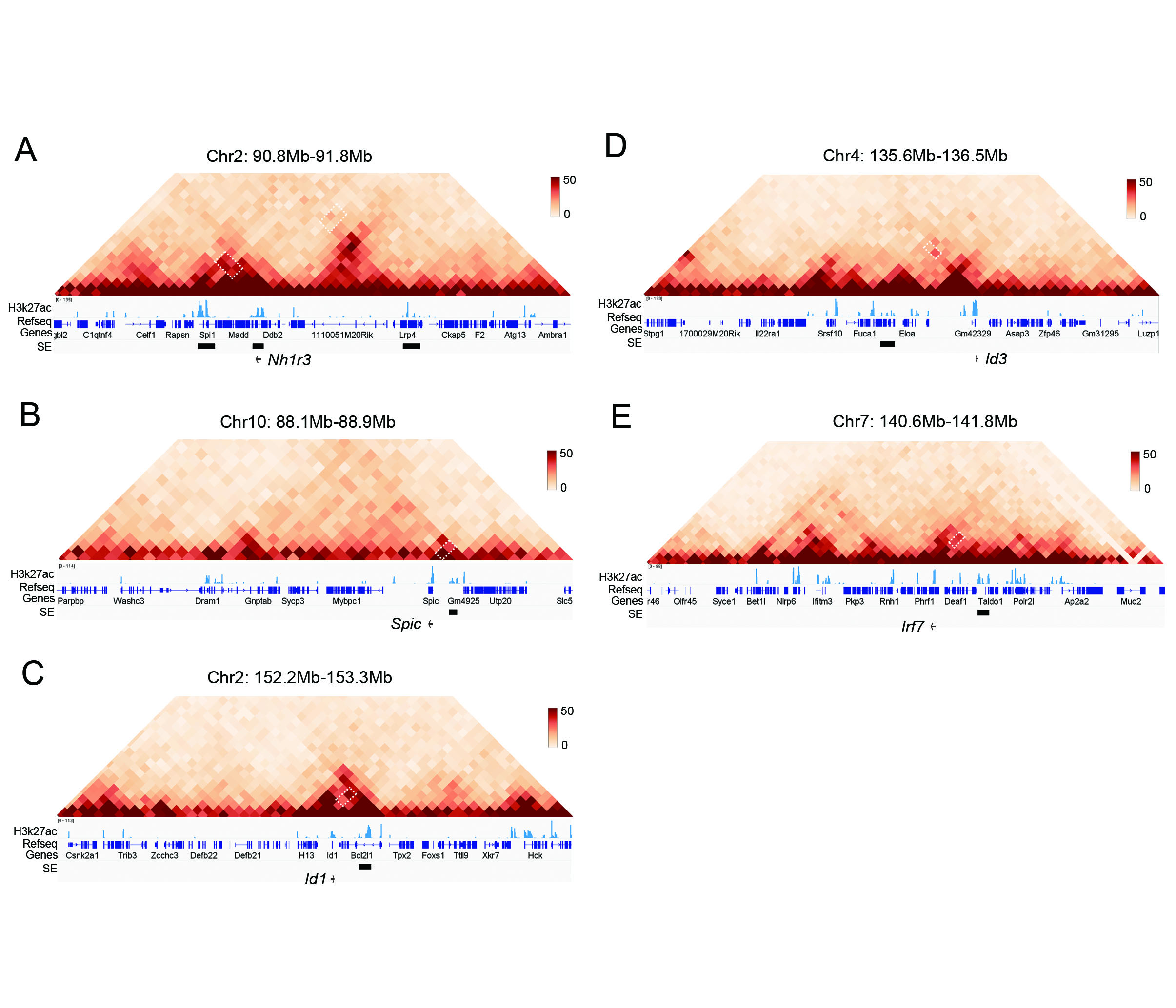
