## Supplemental Table 3 for "Histo-LCM-Hi-C reveals the 3D chromatin conformation of spatially localized rare cells in tissues at high resolution"

**Table S3 Filter results of the sequencing data from published low input Hi-C data**

|  | **2018, Diaz et al.** | **2020, Zhang et al.** | **2020, Zhang et al.** |
| --- | --- | --- | --- |
| Total reads | 86,616,647 | 79,596,960 | 93,469,253 |
| MAPQ≥30 Count | 57,057,214 | 45,191,240 | 71,513,055 |
| Duplicate Removed | 29,559,433 | 34,405,720 | 21,956,198 |
| Nonspecific | 2,749,046 | 9,069,175 | 6,630,422 |
| Valid Pairs | 26,810,387 | 25,336,545 | 15,325,776 |
| Cis-interaction | 22,811,714 | 19,896,310 | 11,764,130 |
| cell number | 1,000 | 500 | 200 |
| Valid/cell | 26,810 | 50,673 | 76,629 |
