## Supplemental Table 4 for "Histo-LCM-Hi-C reveals the 3D chromatin conformation of spatially localized rare cells in tissues at high resolution"

**Table S4 Significant contacts from FitHiC2 in the small-cell-number (KC) datasets before and after down-sampling**

|  | **Before down-sampling** | **After down-sampling** | **Overlap** | **Before down-sampling/**  **overlap%** | **After down-sampling/**  **overlap%** | |
| --- | --- | --- | --- | --- | --- | --- |
| Contacts  (q-value < 0.05) | 4,690,457 | 4,408,583 | 4,310,396 | 91.90% | | 97.77% |
| Contacts  (q-value < 0.01) | 3,011,875 | 2,836,947 | 2,760,925 | 91.67% | | 97.32% |
| Contacts  (q-value < 0.005) | 2,681,144 | 2,512,675 | 2,456,975 | 91.64% | | 97.78% |
